# Distance Mapping and Variable-Specific Geometry of Goal-Relevant Frames in the Retrosplenial Cortex

**DOI:** 10.64898/2026.06.01.729188

**Authors:** Yumin Chen, Xinhang Wei, Lingwei Tang, Haibing Xu

**Affiliations:** Key Laboratory of Mental Health of the Ministry of Education, Guangdong-Hong Kong-Macao Greater Bay Area Center for Brain Science and Brain-Inspired Intelligence, Department of Neurobiology, School of Basic Medical Sciences, Southern Medical University, Guangzhou, 510515, China

**Keywords:** Retrosplenial Cortex, Spatial Navigation, Distance-to-Goal, Mixed Selectivity, Representational Geometry, Reference Frame

## Abstract

Goal-directed navigation requires animals to continuously update their position relative to an unmarked goal. Here, we recorded retrosplenial cortex (RSC) activity in freely moving rats during goal-directed navigation and random foraging. We found that RSC neurons encoded the Euclidean distance to the goal, and that this distance representation was selectively biased toward the goal during navigation. This goal-biased signal could not be explained by non-uniform behavioral sampling alone. Task engagement selectively enhanced allocentric head-direction representations anchored to a landmark cue, whereas egocentric boundary-bearing signals showed no detectable task-related enhancement and no detectable goal-centered spatial organization in this task context. Mixed-selective RSC population activity further exhibited variable-specific separability–smoothness geometry: distance-to-goal showed high local smoothness and decoding performance, whereas egocentric boundary bearing showed stronger macro-scale separability. These task-related spatial representations persisted under reduced visual input, suggesting contributions from memory and self-motion signals. Together, these findings indicate that RSC organizes goal-relevant spatial representations in a task-dependent manner.

## Introduction

Effective navigation relies on the continuous updating and maintenance of an accurate representation of the individual’s position relative to the goal. This process requires the integration of spatial variables encoded in multiple reference frames, including allocentric (world-centered) and egocentric (self-centered) representations[1, 2]. In turn, this process shapes the neural representation of goal-relevant spatial information[3–9].

Retrosplenial cortex (RSC) is a critical brain region involved in spatial navigation[10–15]. Evidence from both rodent and human studies demonstrates that lesions or dysfunctions within RSC lead to significant impairments in spatial navigation, underscoring its essential role in this cognitive process[16–22]. In freely moving rodents, although a subset of RSC neurons exhibits stable spatial representations, their spatial information index is relatively low, and sparse firing patterns similar to hippocampal place cells are rarely observed[23–26]. Additionally, experiments using track-based mazes have shown that certain RSC neurons display spatially symmetric activation patterns, which have been proposed as a novel metric for encoding distance—a feature notably absent in the hippocampus[27]. Collectively, these findings suggest that RSC exhibits a distinct spatial representation pattern compared to the hippocampus, with emerging evidence that it may encode distance-related information rather than precise spatial locations.

RSC exhibits extensive connectivity with key spatial representation regions, including the medial temporal lobe, which encodes allocentric spatial representations, and the posterior parietal cortex, which processes egocentric information[28–32]. Additionally, RSC is connected to thalamic regions involved in head direction processing[33]. These anatomical connections provide RSC with access to multiple sources of spatial representation. Functional studies have further shown that, in addition to having a stable firing rate map of the space, RSC is also capable of processing egocentric boundary information, self-motion, and head direction[23, 24, 34–36]. Based on these findings, RSC is considered to act as a mediator between different brain regions involved in spatial cognition, facilitating the integration of diverse information. Importantly, individual RSC neurons often exhibit mixed selectivity, encoding multiple behavioral variables simultaneously rather than representing a single spatial parameter in isolation[23, 37, 38]. However, mixed selectivity alone does not specify the population-level geometry in which co-encoded variables are represented; recent theoretical work has emphasized that neural population geometry can shape downstream computation[39, 40]. Whether and how this rich functional repertoire is recruited to serve goal-directed navigation, and whether co-encoded variables share a common representational geometry, remain open questions.

Current evidence indicates that RSC neurons are involved in encoding landmark information[9, 41–44], with encoding intensity being augmented during task engagement[41]. Furthermore, in the context of a T-maze spontaneous alternation task, RSC firing patterns are capable of representing the forthcoming goal location as rats approach the choice point[45]. Recent studies have also demonstrated that, following goal-directed navigation, egocentric coding within RSC is enhanced at the vertex corresponding to the goal location[46]. Collectively, these findings suggest that RSC activity can be influenced by task-related states. Despite these insights, how RSC encodes the goal and how its spatial representations are modulated by navigational demands remain unresolved.

In this study, we trained freely moving rats to engage in both goal-directed navigation and random foraging tasks within an open environment while concurrently recording neuronal activity in RSC using tetrode arrays. Our aim was to investigate how RSC represents goal-relevant spatial information during navigation and how these representations are modulated by task demands. Our results reveal that RSC neurons encode the Euclidean distance between the animal and its goal, with this distance signal being selectively biased toward the goal. Although individual neurons conjunctively encoded distance with other behavioral variables, different variables adopted distinct representational geometries at the population level. We further observed that allocentric head direction signals were enhanced during navigation and anchored to the landmark, while egocentric representations remained stable. These spatial representations persisted even under reduced visual input. Collectively, these findings elucidate RSC’s versatile representational role in goal-directed navigation.

## Results

### Identification and Characterization of DTG Cells in RSC

We trained five male Long-Evans rats to perform a goal-directed navigation task in a circular maze (Figure 1A), while recording RSC neuronal activity via a microdrive implanted in the right hemisphere of RSC (Figures 1A and S1A). In this task, rats were required to navigate to a fixed, unmarked location at the start of each trial, where they had to wait for at least two seconds to earn a randomly distributed pellet reward (Figure 1A bottom; see Methods for details). To prevent the rats from developing a dependency on a fixed location, we changed the goal zone between each recording session (Figure S8C). Quantitative behavioral analysis showed that rats exhibited more concentrated trajectories around the goal area and reduced their speed when approaching the goal (Figures 1B and S8A). This suggests that the rats were able to recognize the invisible goal zone. On average, the rats completed three trials per minute (mean ± SEM rewards per minute: 3.1 ± 0.8, n = 5 rats, Figure 1C), further supporting their ability to navigate successfully.

**Figure 1.**
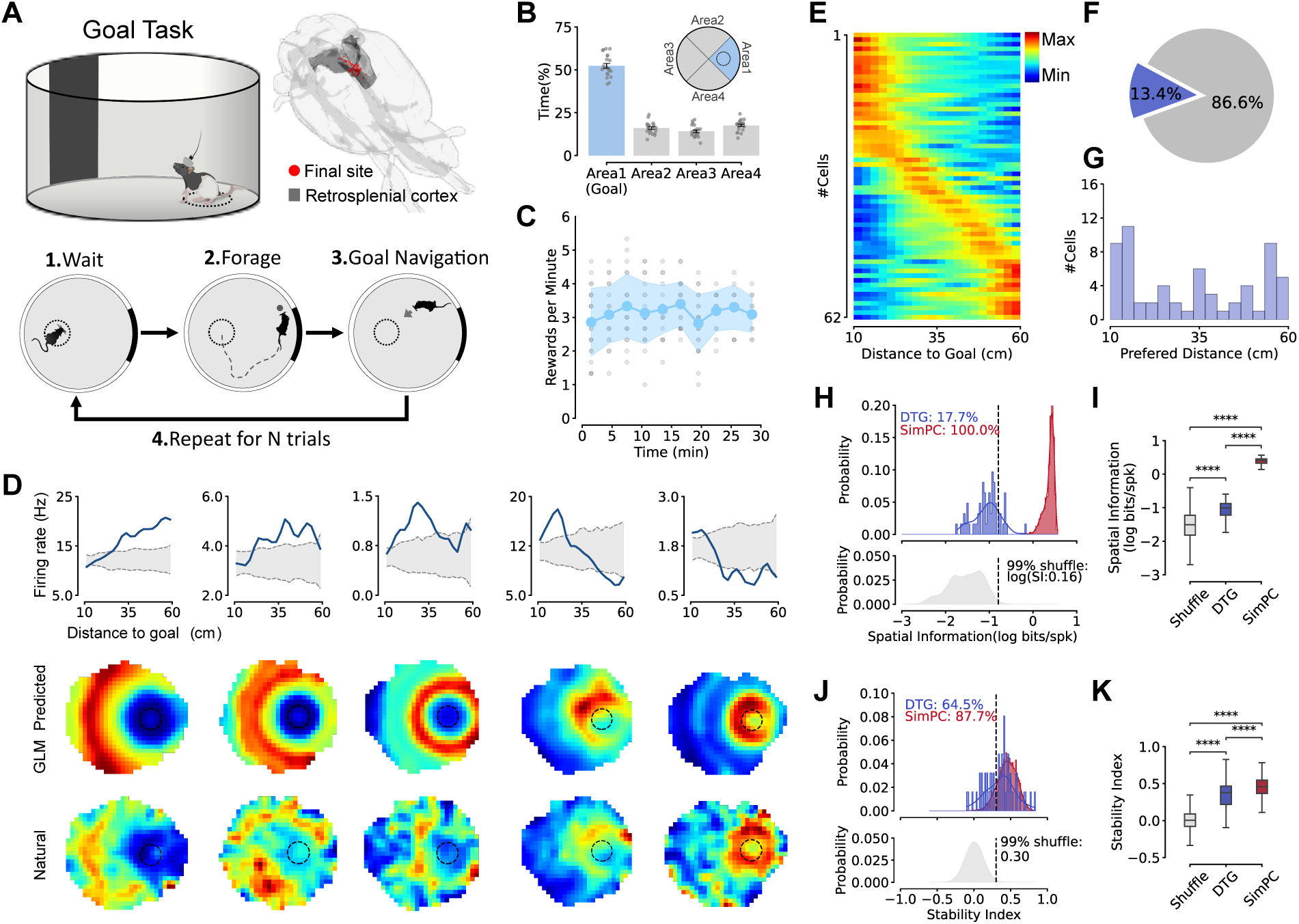
Functional profiling of DTG cells in a goal-directed navigation task. (A) Schematic of the goal-directed navigation task. Top Left: the circular maze with a black cue and an unmarked goal zone (dashed line). Top Right: Reconstructions were made of the final tetrode recording sites (red dots) in the right hemisphere of RSC (dark gray) for all animals. Bottom: Task protocol. The rat is required to wait for at least two seconds in the goal zone, then leave the goal zone to search for a single randomly scattered pellet reward, consume the reward, and subsequently navigate back to the goal zone for the next trial. (B) The maze was evenly divided into four areas, and the trajectory occupancy was calculated for each. The rats’ trajectories were concentrated in the area with goal (blue). The grey dots represent the sessions and the error bars represent mean ± SEM of the data. (C) The reward acquisition across each session is shown, with gray dots representing the number of rewards earned per minute for all sessions at each time point. Blue dots indicate the mean reward rate, and the blue shaded region represents mean ± SD of the reward rate across sessions. (D) Examples of five DTG cells. Top: Distance-to-Goal tuning curves (gray area represents95% CI from shuffle). Middle: GLM-predicted spatial rate maps reconstructed from the fitted Poisson GLM for each cell. Bottom: observed (natural) spatial rate maps. The concordance between GLM-predicted and observed maps confirms that the model captures each cell’s spatial firing structure. (E) Sorted tuning curves of all DTG cells (n = 62). (F) Proportion of cells encoding Distance-to-Goal identified through the intersection ofGLM-based forward feature selection and shuffle-based significance criteria (DTG: 13.4%, 62/463). (G) Distribution of preferred distances for DTG cells. (H) Spatial information index distribution of DTG cells. Top: spatial information indexhistogram for DTG cells (blue) and simulated place cells (red). Bottom: null distribution from shuffle group; black dashed line indicates the 99% threshold from the shuffle group. (I) Comparison of the spatial information index among three groups. DTG cells have significantly higher spatial information index values than the shuffle group, but their spatial information index values are significantly lower than those of simulated place cells. **p < 0.0001. (J) Similar to panel H but showing spatial stability. Top: Stability histogram for DTG cells (blue, 64.5% above threshold) and simulated place cells (red, 87.7% above threshold). Bottom: null distribution from shuffle group; black dashed line indicates the 99% threshold (stability = 0.30) from the shuffle group. (K) Comparison of spatial stability among three groups. DTG cells have significantly higherstability than the shuffle group, but are significantly lower than simulated place cells. **p < 0.0001.

A total of 463 RSC neurons were analyzed in the goal-directed navigation task, including 88.6% putative pyramidal neurons (Pyr) (410/463, 5 rats, 21 sessions) and 11.4% interneurons (Int) (53/463). Of the total population, 68.3% (n = 316/463) were recorded from dysgranular RSC (dRSC), and 31.7% (147/463) within granular RSC (gRSC) (Figures S1). Using a GLM-based forward feature selection combined with a shuffle-based significance procedure and stability quantification (see Methods), we identified a group of cells that can tune to the Euclidean distance between the animal and its goal (13.4%, 62/463 from 5 rats, Pyr: 93.5%, 58/62, Int: 6.5%, 4/62; dRSC: 72.6%, 45/62, gRSC: 27.4%, 17/62, Figures 1E, 1F and S1D). We designate this group of neurons that encode Distance-to-Goal information as DTG cells. The spatial firing rate maps of most DTG cells exhibit a ring-like firing pattern centered around the goal (Figures 1D middle and S2A). The preferred distances of DTG cells spanned a broad range (Figure 1G), and cells tuned to short distances also displayed ring-like spatial rate maps centered on the goal, consistent with the overall population pattern (Figure 1D).

It is not difficult to conclude that DTG cells are not necessarily place cells. However, place cells can be considered as DTG cells, as a specific place representation can also be interpreted as a distance representation to a particular point. To further characterize the spatial firing properties of DTG cells, we quantified and compared the spatial firing properties of DTG cells, other RSC cells, and simulated place cells (Figures 1H-1K and S2). DTG cells exhibited significantly higher spatial information index than the shuffle group, but were significantly lower than simulated place cells (Wilcoxon rank-sum test for Shuffle vs. DTG, U=-9.14, p<0.0001; for Shuffle vs. SimPC, U=-66.45, p<0.0001; for DTG vs. SimPC, U=-13.36, p<0.0001; Figures 1H, 1I, and S2C). Similarly, DTG cells showed significantly higher spatial stability than the shuffle group, but were significantly lower than simulated place cells (Wilcoxon rank-sum test for Shuffle vs. DTG, U=-11.70, p<0.0001; for Shuffle vs. SimPC, U=-65.31, p<0.0001; for DTG vs. SimPC, U=-4.34, p<0.0001; Figures 1J, 1K and S2D). These results solidify the RSC’s unique computational profile: DTG cells exhibit enhanced spatial properties relative to the general RSC population, yet remain below simulated place cells in both spatial information and stability, consistent with a prioritization of distance coding over high spatial information, underscoring the RSC’s specialized role in translating dynamic spatial metrics into goal-directed navigation.

Although individual DTG cells clearly encoded distance-to-goal, their spatial information and stability remained below those of simulated place cells, suggesting that the full distance representation in RSC may rely on the coordinated activity of a neural population rather than on a small number of highly informative single neurons. We therefore asked how the RSC neural population collectively encodes Distance-to-Goal, and whether a specific subset of neurons disproportionately contributes to this representation.

### Sparse Efficient Encoding of Distance-to-Goal in RSC Neural Population

To address this question, we applied CEBRA, a self-supervised nonlinear dimensionality reduction method, to obtain low-dimensional latent representations of RSC population activity and performed supervised decoding of Distance-to-Goal on sessions containing at least 20 simultaneously recorded cells (n = 9 sessions, from 5 animals; Figure 2A). The population decoder faithfully tracked the animal’s moment-to-moment distance to the goal, confirming that Distance-to-Goal is robustly represented at the population level in RSC (Figure 2A).

**Figure 2.**
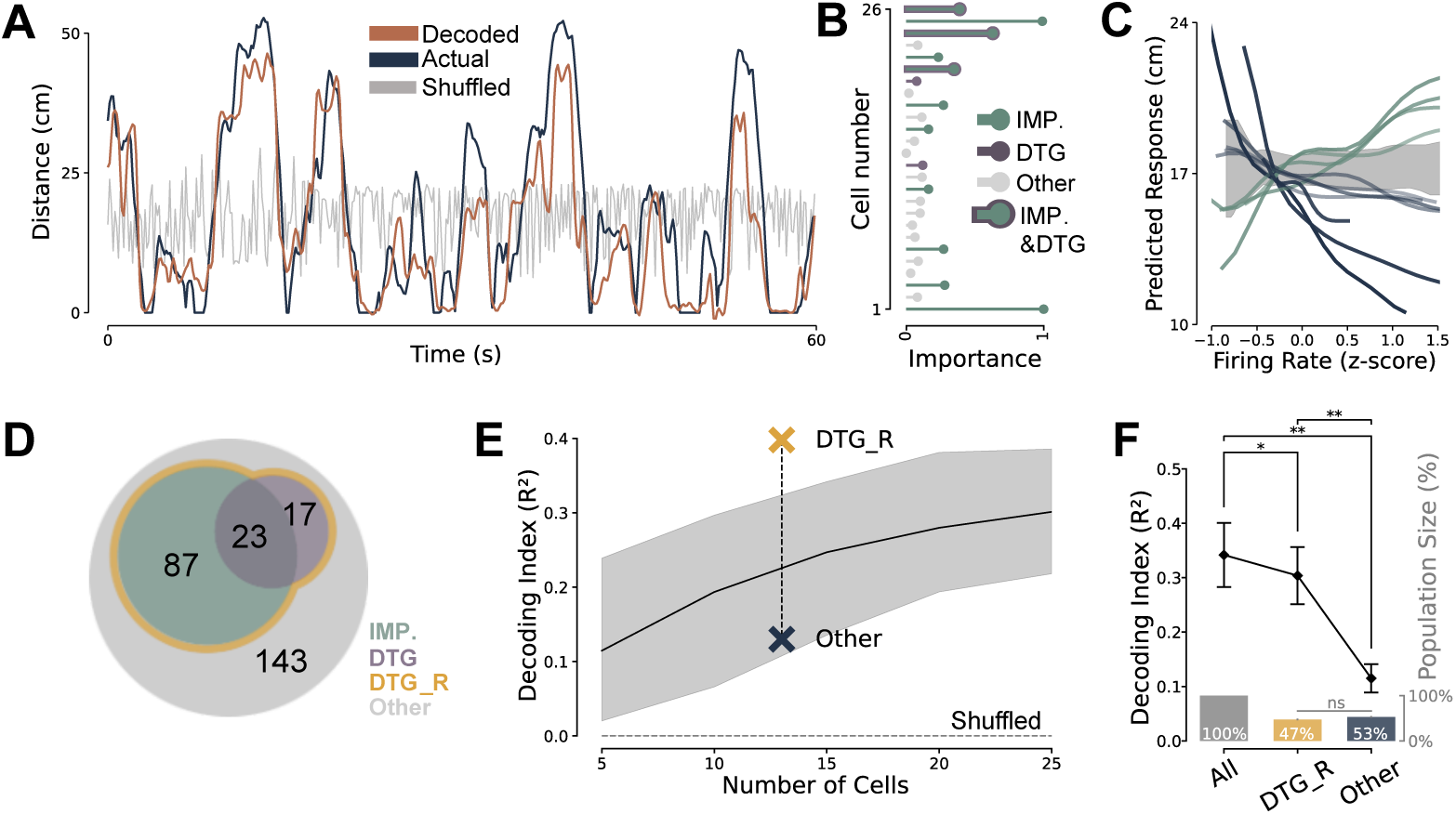
Sparse efficient encoding of Distance-to-Goal in RSC neural population. (A) Comparison of decoded (orange), actual (dark blue), and shuffled (grey) Distance-to-Goal values over a representative 60-second segment, demonstrating that the CEBRA-based population decoder faithfully tracks the animal’s moment-to-moment distance to the goal. (B) Evaluation of the contribution of single neurons in decoding Distance-to-Goal, usingimportance score calculated from EBM. IMP (green), DTG (purple), IMP & DTG (green & purple), and Other (grey). Each bar represents the importance score of a cell, with the length of the bar indicating the magnitude of the score. (C) Partial dependence plot (PDP) of predicted response versus firing rate (z-scored) for eachIMP cell. IMP cells are shown as individual lines with color-coding based on the sign of their linear slope (green for positive, dark blue for negative). Other neurons are represented as shaded grey areas, providing an overview of general response patterns. Both IMP and DTG cells exhibit monotonic relationships between firing rate and predicted Distance-to-Goal, whereas Other RSC cells display flat, uninformative response profiles. (D) Venn diagram showing the overlap between different neuron groups (IMP only = 87,DTG only = 17, IMP *∩* DTG = 23, DTG_R = 127 total, Other = 143). (E) Effect of cell number on Distance-to-Goal decoding performance. The black line representsthe average decoding performance, with the gray shaded region showing the 95% CI from 100 sampling iterations. The gray dashed line represents the shuffled baseline results. The DTG_R group (golden cross) exceeds the expected decoding performance for its population size, while the Other group (dark cross) shows the lowest decoding performance for its population size. (F) Decoding index (*R*^2^) across different cell groups. DTG_R cells exhibit significantly higher decoding performance compared to the Other group, and All cells significantly outperform DTG_R (Wilcoxon signed-rank test: ALL vs. DTG_R, W = 5, p = 0.039; ALL vs. Other, W = 0, p = 0.004; DTG_R vs. Other, W = 0, p = 0.004; n = 9 sessions, two-sided). Diamond point and error bars represent mean ± SEM of the data. *p < 0.05, **p < 0.01. Inset: population sizes of All, DTG_R, and Other groups as percentage of total.

A critical question arising from the population decoding is whether all RSC neurons contribute equally to this representation, or whether a smaller subset carries the bulk of the signal. To dissect this, we applied the Explainable Boosting Machine (EBM) model to the same population activity, utilizing its interpretability to pinpoint the key neural population involved in Distance-to-Goal encoding. Based on the importance scores provided by the EBM and the single-cell DTG classification (see Methods), cells are categorized into four groups: IMP cells (identified as strongly involved in decoding based on EBM importance scores), DTG cells (identified through the intersection of single-cell Distance-to-Goal tuning and GLM-based selection criteria), IMP & DTG cells, and Other RSC cells (Figures 2B and 2C). Although IMP cells exhibited slightly lower DTG scores compared to DTG cells, their scores were still significantly higher than those of other RSC cells. Besides, the Partial dependence plots (PDPs) of the EBM model showed that both IMP and DTG cells exhibit monotonic relationships between firing rate and predicted Distance-to-Goal with individual neurons showing either positive or negative slopes whereas Other RSC cells display flat, uninformative response profiles (Figures 2C and S3).

Based on these findings, the IMP and DTG cell populations were combined into a group termed DTG-relative cells (DTG_R), which is considered the subgroup most closely associated with Distance-to-Goal encoding (IMP only=87, DTG only=17, IMP*∩*DTG=23, DTG_R=127 total, Other=143; Figure 2D). Notably, the DTG_R group comprised 47.0% of neurons included in the population-decoding sessions (127/270), yet captured a disproportionately large share of the Distance-to-Goal signal. The decoding results confirmed this sparse efficient encoding: despite the general trend that more cells yield better accuracy (Figure 2E), DTG_R cells achieved decoding performance approaching that of the full population, while significantly outperforming the Other cells (Distance-to-Goal decoding index: Wilcoxon signed-rank test for ALL vs. DTG_R, W=5, p=0.039; for ALL vs. Other, W=0, p=0.004; for DTG_R vs.

Other, W=0, p=0.004; n = 9 sessions, two-sided; Figure 2F). In other words, nearly half of the neurons included in population decoding contributed minimally to distance decoding, while the DTG_R subset alone accounted for the majority of the population’s representational capacity. The convergence of two independent identification approaches bottom-up singlecell tuning analysis (DTG) and top-down population-level importance scoring (IMP) on a substantially overlapping set of neurons (Figure 2D) provides cross-validated evidence that Distance-to-Goal encoding in RSC is concentrated in a functionally coherent subpopulation.

The preceding analyses establish that RSC contains a functionally enriched subpopulation for Distance-to-Goal encoding, but they do not address whether this representation is spatially uniform across the environment or preferentially directed toward the goal. Moreover, because the animals’ trajectories are inherently non-uniform during goal-directed navigation, concentrating around the goal area (Figure 1B), it is essential to determine whether the observed distance signals reflect neural encoding beyond behavioral sampling biases or merely mirror those biases. We next addressed both questions by constructing spatial attention maps and introducing a simulation-based control.

### Task-Dependent Goal-Biased Distance Encoding in RSC Population

To characterize the spatial focus of the RSC distance representation, we calculated the population-level Distance-to-Goal decoding performance for various points in space and constructed an attention map representing the animal’s spatial preference. The attention map allows us to visually identify the focal point of the RSC neuronal distance representation and assess whether there is a consistent positional encoding preference (Figure 3A).

**Figure 3.**
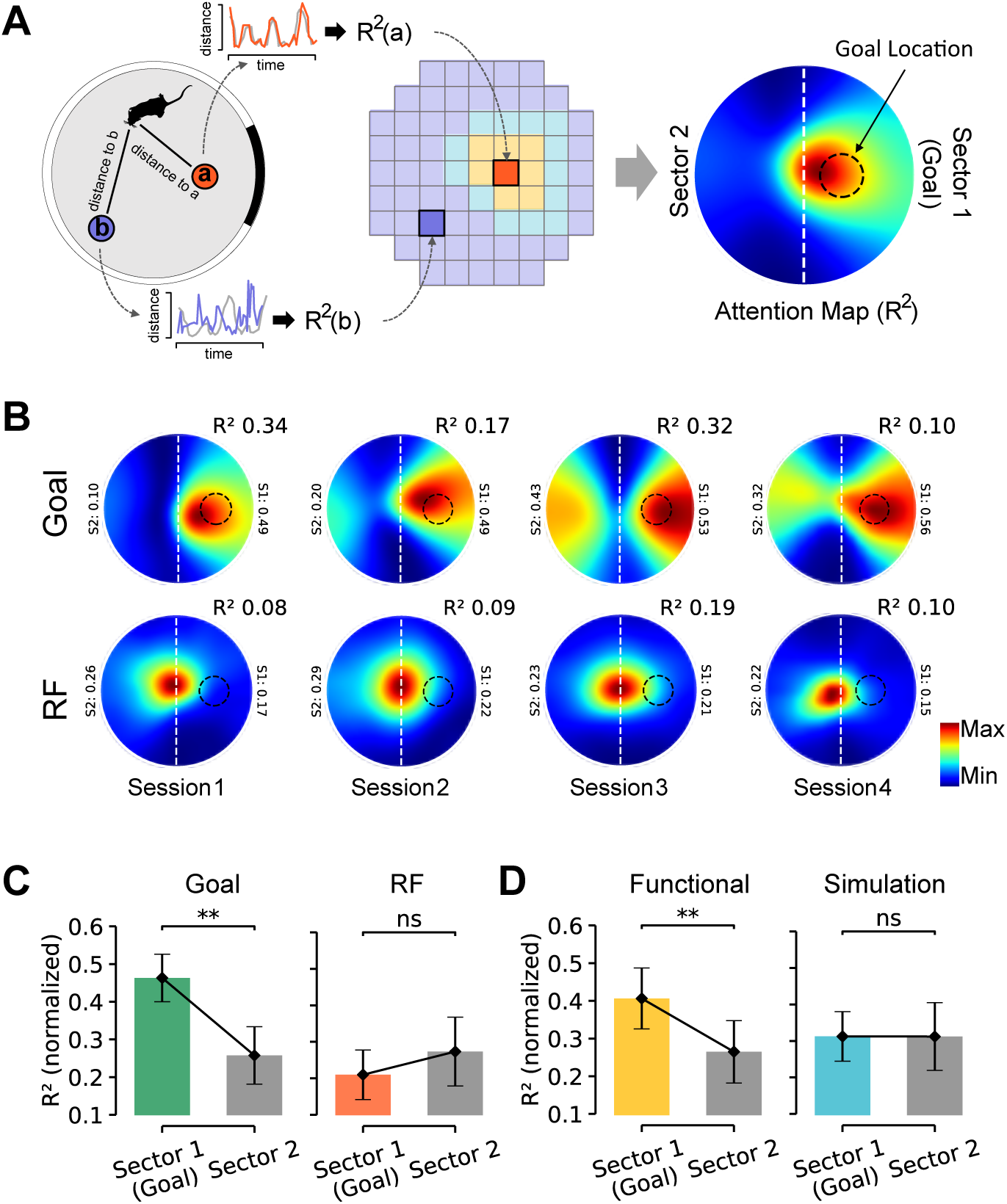
Task-dependent goal-biased distance encoding revealed by neural attention map. (A) A schematic of attention map calculation. Left: For each reference point in the maze, thedistance from the rat to that point is calculated and decoded, with the resulting *R*^2^ value used to quantify the animal’s representation strength for that location. Middle: These points are used to construct an attention map that represents the spatial attention distribution of the animal regarding distance encoding. Right: The final attention map is divided into two sectors based on the location of the Goal: Sector 1 (Goal) and Sector 2. The mean normalized *R*^2^ values in both sectors are then calculated and used to analyze the relationship between the animal’s distance representation and the Goal. (B) Attention maps from the goal-directed navigation task and the corresponding maps from the random foraging task for four example sessions. (C) Left: In the goal-directed navigation task, *R*^2^ values are significantly higher in Sector 1 (with the Goal) compared to Sector 2 (Wilcoxon signed-rank test, n = 9, p = 0.002). Right: In the random foraging task, no significant difference is found in attention maps between the two sides of the maze (Wilcoxon signed-rank test, n = 8, p = 0.991). p < 0.01, ns > 0.05. (D) Similar to C. Compared to the functional population’s preference coding for the goal area(Wilcoxon signed-rank test, n = 9, p = 0.002), simulated cells show no preference for the goal side in their attention maps (Wilcoxon signed-rank test, n = 9, p = 0.633). p < 0.01, ns > 0.05.

We found that, during the goal-directed navigation task, the neural population’s distance representation exhibited a clear preference for the goal side (Goal-Light: Wilcoxon signed-rank test for Sector 1 vs. Sector 2, W=45, p=0.002, n=9 sessions, one-sided; Figures 3B top and 3C left). In contrast, during the random foraging task, the attention focus of the distance representation was located at the center of the maze (RF-Light: Wilcoxon signed-rank test for Sector 1 vs. Sector 2, W=0, p=0.991, n=8 sessions, one-sided; Figures 3B bottom and 3C right). We attribute this to a preference coding of the boundary (or maze center, which is computationally equivalent).

A potential confound is that goal-directed behavior produces non-uniform spatial sampling, which could artifactually inflate decoding near the goal. To control for this, we constructed a simulated neural population using RatInABox[47], in which each recorded neuron was replaced by a simulated counterpart that faithfully reproduced its known functional tuning property (e.g., head direction, egocentric boundary bearing, or speed) but contained no explicit Distance-to-Goal signal (Figure S4A–E). Critically, these simulated neurons were driven by the animal’s real trajectory, thereby inheriting the same behavioral non-uniformities as the recorded population. The resulting simulated cells closely matched their real counterparts in tuning similarity (Figure S4F). Despite this high fidelity in replicating known functional properties, the simulated population exhibited substantially weaker Distance-to-Goal tuning: functional cells showed significantly steeper Distance-to-Goal tuning slopes than their simulated counterparts (Figure S4G), and the functional population significantly outperformed the simulated population in Distance-to-Goal decoding (Wilcoxon signed-rank test for Data vs. Sim., W=1, p=0.027, n=9 sessions, two-sided; Figure S4H). These results indicate that the simulation effectively isolates the contribution of the tested behavioral biases from the recorded neural distance signal.

We then applied the same attention map analysis to functional cells and their corresponding simulated cells. The results showed that only the functional cell population exhibited a goal-preference distance representation despite both populations containing the same influence from goal-directed non-uniform trajectory (Functional: Wilcoxon signed-rank test for Sector 1 vs. Sector 2, W=45, p=0.002; Simulated: Wilcoxon signed-rank test for Sector 1 vs. Sector 2, W=20, p=0.633; n=9 sessions, one-sided; Figures 3D and S5). These findings demonstrate that the goal-biased distance representation in RSC cannot be explained by non-uniform behavioral sampling alone, but instead reflects a neural population bias not attributable to the simulated behavioral confounds tested here. Combined with the sparse efficient encoding identified above, these results indicate that RSC shows a task-dependent bias toward the goal as the reference point for distance encoding.

### Mixed Selectivity and Variable-Specific Representation Geometry in RSC

The analyses above establish that RSC contains a functionally coherent subpopulation enriched for Distance-to-Goal encoding, and that this representation is biased toward the goal during navigation. However, they treat Distance-to-Goal in isolation. In practice, RSC neurons are known to encode multiple behavioral variables simultaneously[23, 24, 34, 37, 48]. We therefore first asked whether DTG encoding was represented in isolation or embedded within the broader mixed-selectivity landscape of RSC.

To characterize mixed selectivity at the single-cell level, we applied a Poisson GLM with forward feature selection to quantify the contribution of five behavioral variables head direction (H), speed (S), distance-to-goal (D), egocentric boundary bearing (E), and position (P) to the firing of each neuron (Figure 4A; see Methods). Because the task structure can introduce correlations between behavioral variables, such as the coupling between distance-to-goal and speed as animals decelerate near the goal, L1 regularization was used to ensure that each retained variable made an independent contribution to firing-rate prediction. The majority of DTG cells encoded multiple variables conjunctively: 35.5% (22/62) encoded two variables, 17.7% (11/62) encoded three, and 22.6% (14/62) encoded four, while 24.2% (15/62) encoded a single variable (Figure 4B). An example DTG cell illustrating conjunctive tuning to head direction, speed, and egocentric boundary bearing is shown in Figure 4C.

**Figure 4.**
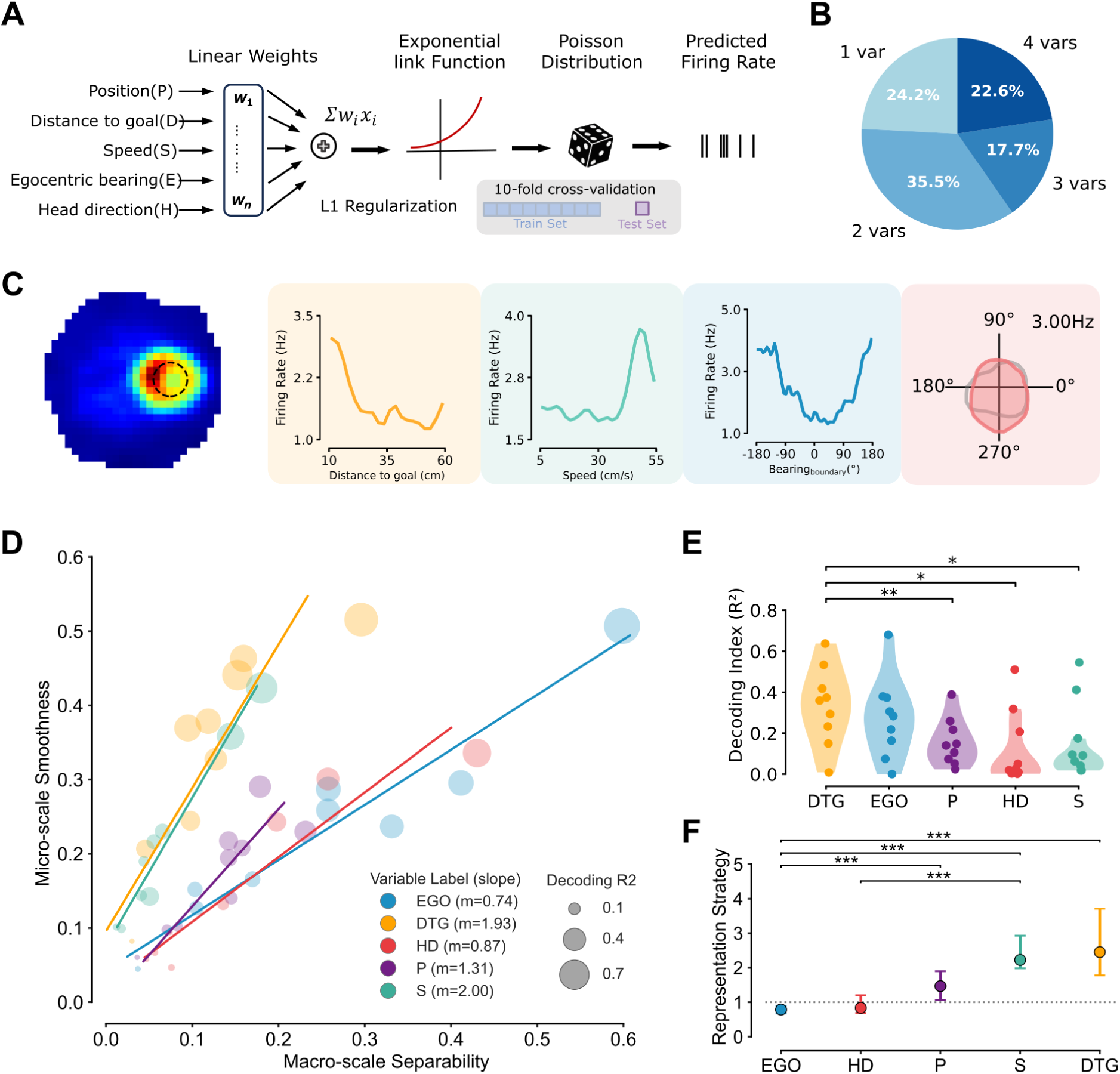
Mixed selectivity and variable-specific representation geometry in RSC. Panels A–C characterize mixed selectivity of DTG cells, whereas panels D–F quantify variable-specific separability–smoothness geometry at the population level. (A) Schematic of the Poisson GLM with forward feature selection. Five behavioral variablesPosition (P), Distance-to-Goal (D), Speed (S), Egocentric boundary bearing (E), and Head direction (H) are each projected onto B-spline basis functions and combined through linear weights with L1 regularization. The weighted sum is passed through an exponential link function to produce Poisson-distributed predicted firing rates. Model selection is performed via 10-fold cross-validation with a forward search procedure (see Methods). (B) Distribution of the number of behavioral variables encoded by DTG cells as determined bythe GLM forward search. The majority of DTG cells encode multiple variables conjunctively: 35.5% (22/62) encode two variables, 17.7% (11/62) encode three, and 22.6% (14/62) encode four, while 24.2% (15/62) encode a single variable. (C) An example DTG cell illustrating conjunctive encoding. Left: spatial rate map (GLMpredicted) showing a ring-like firing pattern centered on the goal. Right: tuning curves for Distance-to-Goal, Speed, and Egocentric boundary bearing, and a polar plot for Head direction, demonstrating that this cell simultaneously encodes multiple behavioral variables. (D) Separability–smoothness geometry for GoalLight sessions. Each variable is summarizedby macro-scale separability (structure index, StrI; x-axis) and Micro-scale Smoothness (1*−* normalized Local Neighborhood Consistency; y-axis). Dot size indicates decoding *R*^2^. Positive slopes describe the balance between separability and local smoothness for each variable: EGO (blue, *m* = 0.74), DTG (orange, *m* = 1.93), HD (red, *m* = 0.87), P (purple, *m* = 1.31), and S (teal, *m* = 2.00). (E) Decoding Index (*R*^2^) comparison across five behavioral variables. DTG and EGO achieve the highest decoding performance, while adopting opposite representation geometries (see D). *p < 0.05, **p < 0.01. (F) Bootstrap stability analysis of separability–smoothness slopes across GoalLight andGoalDark sessions. Points and intervals show mean bootstrap slope estimates and 95% confidence intervals (n = 10,000 iterations). The dashed line marks the unity reference (*m* = 1). Pairwise significance markers indicate permutation-test differences in slope between variables. **p < 0.01, ***p < 0.001.

We next examined whether any specific functional cell type was preferentially represented within the DTG population. The proportions of HD cells, EGO cells, speed cells, and place-encoding cells among DTG neurons did not differ significantly from those in the overall population (HD: All = 16.4%, DTG = 19.4%, p = 0.63; EGO: All = 17.3%, DTG = 17.7%, p = 1.0; Speed: All = 19.2%, DTG = 12.9%, p = 0.24; Place: All = 19.9%, DTG = 19.4%, p = 1.0; Chi-squared test; Figure S6). Thus, Figure 4A–C indicates that DTG cells participate in broad mixed selectivity rather than forming a narrowly isolated distance-only class.

Having established mixed selectivity at the single-cell level, we next asked whether the co-encoded variables were organized similarly or differently at the population level. To address this question, we quantified each variable in a two-dimensional geometry space combining macro-scale separability and micro-scale smoothness (Figure 4D). Macro-scale separability was measured by the structure index (StrI), which quantifies how well samples with similar labels are organized at the population level. Micro-scale smoothness was defined as 1*−* normalized Local Neighborhood Consistency (LNC), so that larger values indicate more locally coherent label organization in the embedding.

In GoalLight sessions, Distance-to-Goal (DTG) showed the highest micro-scale smoothness and decoding performance among the examined variables (mean smoothness = 0.337; mean decoding *R*^2^ = 0.328), whereas egocentric boundary bearing (EGO) showed the highest macro-scale separability (mean StrI = 0.252). Other variables occupied lower or intermediate positions in this separability–smoothness space, including Head direction (HD; mean StrI = 0.157, mean smoothness = 0.158), Position (P; mean StrI = 0.132, mean smoothness = 0.171), and Speed (S; mean StrI = 0.068, mean smoothness = 0.212). Thus, mixed selectivity in RSC was accompanied by variable-specific population geometry rather than a single shared embedding profile across all encoded variables.

We next summarized the balance between separability and micro-scale smoothness using the slope of each variable’s trajectory in separability–smoothness space. In GoalLight sessions, DTG and S showed steeper smoothness-weighted slopes (DTG, *m* = 1.93; S, *m* = 2.00), P was intermediate (*m* = 1.31), and EGO and HD showed lower slopes (EGO, *m* = 0.74; HD, *m* = 0.87; Figure 4D). A bootstrap stability analysis pooling GoalLight and GoalDark sessions confirmed this ordering at the group level: EGO and HD remained below or near the unity reference on average, whereas P, S, and DTG showed slopes above unity (Figure 4F). Pairwise permutation tests identified significant slope differences for EGO versus DTG, EGO versus S, EGO versus P, and HD versus S. Together, Figure 4 shows two related but distinct points: RSC neurons often multiplex distance-to-goal with other task variables, and these co-encoded variables retain variable-specific population geometries. The slope metric is used here as an operational descriptor of population geometry, not as evidence for a discrete coding mechanism, an optimality principle, or a downstream readout mechanism.

### Allocentric Enhancement and Stable Egocentric Representations in RSC during Goal-Directed Navigation

The preceding analyses characterize RSC’s representational content and geometry during goal-directed navigation. We next asked how task demands modulate the strength of these co-existing spatial representations by comparing neuronal activity between goal-directed navigation and random foraging. To compare the impact of navigational demands on RSC, we ran random foraging task on the day of or the day before the goal-directed navigation task (see Methods for details). In the random foraging task, we recorded a total of 464 RSC neurons, with 7.5% (35/464, 5 rats, 20 sessions) identified as HD cells, a proportion significantly lower than the 16.4% (76/463) HD cells observed in the goal-directed navigation task (*χ*^2^ = 17.30, p = 3.18 *×* 10*^−^*^5^; with Chi-squared test; Figures 5A and 5B top). And decoding analysis revealed that the decoding accuracy of the head-direction in the goal-directed navigation task was significantly higher than in the random foraging task (Head-direction decoding index: Wilcoxon rank-sum test for Goal vs. RF, U=2, p=0.043; Goal=9 sessions, RF=8 sessions; Figure 5B bottom).

**Figure 5.**
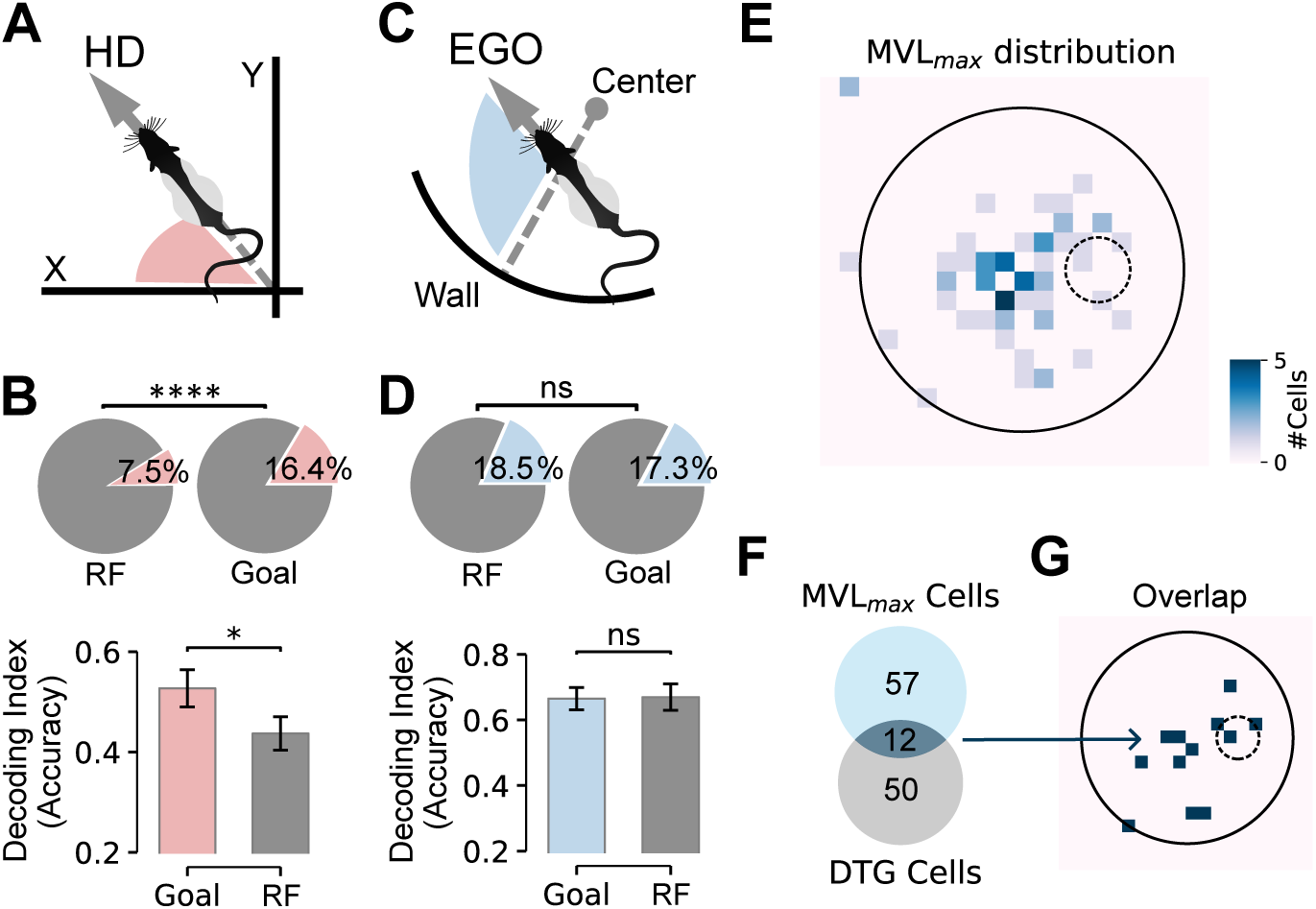
Allocentric enhancement and stable egocentric representations in RSC during goal-directed navigation. (A) Schematic of head direction (red) calculation. (B) Top: the proportion of head-direction cells in the goal-directed navigation task vs. thatin the random foraging task (RF: 7.5%, 35/464; Goal: 16.4%, 76/463; *χ*^2^ = 17.30, p = 3.18 *×* 10*^−^*^5^). Bottom: decoding index (accuracy) of head direction using all neurons in the goal-directed task vs. that in the random foraging task (Wilcoxon rank-sum test, U = 2, p = 0.043; Goal = 9 sessions, RF = 8 sessions). ****p < 0.0001, *p < 0.05. (C) Schematic of Egocentric-boundary-bearing (blue) calculation. (D) Top: the proportion of egocentric boundary bearing cells in the goal-directed navigationtask vs. that in the random foraging task (RF: 18.5%, 86/464; Goal: 17.3%, 80/463; *χ*^2^ = 0.25, p = 0.618). Bottom: decoding index (accuracy) of Egocentric-boundary-bearing using all neurons in the goal-directed navigation task vs. that in the random foraging task (Wilcoxon rank-sum test, U = −0.4, p = 0.7; Goal = 9 sessions, RF = 8 sessions). ns > 0.05. (E) Spatial distribution of MVL_max locations for neurons with high egocentric tuningselectivity (MVL_max > 0.25). The heatmap shows the number of cells whose MVL_max falls in each spatial bin. Neurons with high MVL_max are distributed across the maze without specific concentration near the goal region (dashed circle), in contrast to the goal-biased pattern observed in the population-level distance representation (Figure 3B). (F) Venn diagram showing the overlap between MVL_max neurons and DTG neurons (MVL_max only = 57, overlap = 12, DTG only = 50), confirming that these represent largely distinct subpopulations. (G) Spatial distribution of MVL_max locations for the 12 co-classified neurons (overlapbetween MVL_max and DTG populations). These neurons show no preferential MVL_max locations near the goal (dashed circle), indicating that egocentric boundary coding showed no detectable goal-centered spatial organization in this task context.

In contrast to the head-direction, which is considered an allocentric type of spatial representation, we did not observe a strengthening of egocentric representations during the goal-directed navigation task. The number of EGO cells and the decoding accuracy of Egocentric-boundary-bearing were similar between the goal-directed navigation task and the random foraging task (EGO cells in RF = 18.5% (86/464), EGO cells in Goal = 17.3% (80/463), *χ*^2^ = 0.25, p = 0.618, ns; with Chi-squared test; Egocentric-boundary-bearing decoding index: Wilcoxon rank-sum test for Goal vs. RF, U=-0.4, p=0.7; Goal=9 sessions, RF=8 sessions; Figures 5C and 5D).

To further examine whether egocentric coding contributes to goal-specific spatial representation, we followed the method proposed by Wang et al.[9] and calculated the MVL of the egocentric bearing tuning for each RSC neuron at every point in space, identifying the spatial location where each neuron’s egocentric tuning was strongest (MVL_max). We found that neurons with high MVL_max (>0.25) were distributed across the maze without specific concentration near the goal region (Figure 5E), in contrast to the goal-biased pattern observed in the population-level distance representation (Figure 3B). Moreover, the overlap between MVL_max neurons and DTG neurons was minimal (MVL_max only = 57, DTG only = 50, overlap = 12; Figure 5F), confirming that these represent largely distinct subpopulations. The 12 co-classified neurons showed no preferential MVL_max locations near the goal (Figure 5G). Based on these findings, we conclude that goal-directed navigation selectively enhances allocentric head-direction representations in RSC, whereas egocentric boundary coding showed no detectable task-related enhancement and no detectable goal-centered spatial organization in this task context.

### Stability of Goal Encoding in RSC during Dark Navigation

Because the goal position varied across sessions while the cue remained fixed (Figure S8C), the angular relationship between these two references differed from session to session, allowing us to dissociate whether HD cells are preferentially anchored to the stable visual landmark or to the variable goal location. To this end, we defined two angular coordinate systems: the cue-aligned system, which sets the direction of the landmark cue as 0 degrees, and the goal-aligned system, which sets the direction of the goal as 0 degrees (Figure S7). In the cue-aligned coordinate system, we calculated the angular preference distribution of all HD cells. Our results showed that in the goal-directed navigation task, HD cells exhibited a clear preference for the cue (Rayleigh p=0.004, n=95 cells; Figure 6A middle). In the random-foraging task, HD preferred angles did not show significant cue-aligned clustering (Rayleigh p = 0.472, n = 39 cells; Figure 6A top). Interestingly, using the goal-aligned angular coordinate system, the preferred angles of HD cells did not show significant goal-aligned clustering (Figures S7).

**Figure 6.**
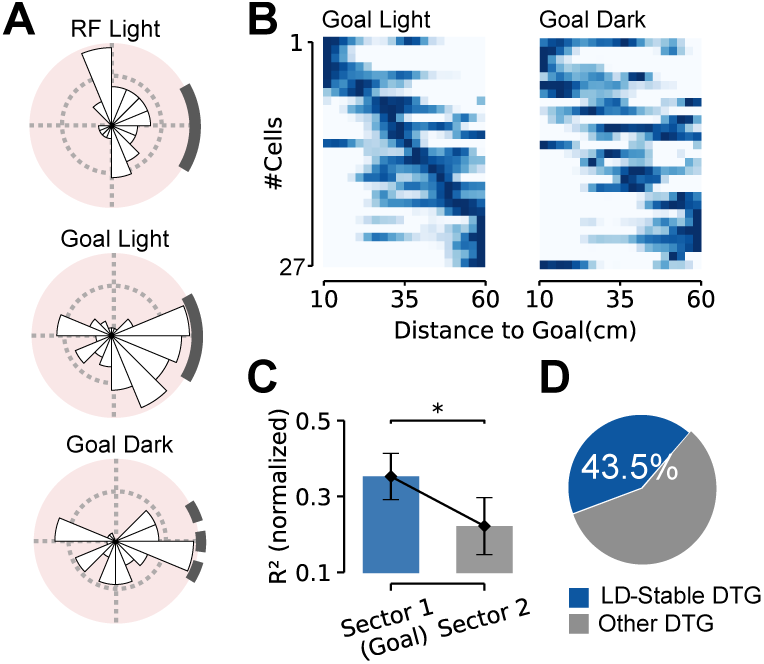
Stability of goal encoding in RSC during dark navigation. (A) Preferred angle of HD cells across three behavioral tasks (RF Light, Goal Light, GoalDark). The cue is positioned on the right side of each figure (dark gray). Note that in the dark environment, the cue will be removed; the original position is shown here. In the goal-directed navigation task, HD cells exhibited a clear preference for the cue (Rayleigh p = 0.004, n = 95 cells). In the random foraging task, no preference for the cue was observed (Rayleigh p = 0.472, n = 39 cells). In the dark condition, HD cells still maintained a preference for the direction of the original cue (Rayleigh p = 0.01, n = 71 cells). (B) Tuning curves of all DTG cells that maintained Distance-to-Goal tuning across light and dark conditions (n = 27 cells), displayed as population heatmaps. Left: Goal Light condition. Right: Goal Dark condition. Cells are sorted by the position of the peak tuning value in the light condition, demonstrating preservation of tuning order across conditions. (C) In the dark environment, the distance encoding of RSC neurons still preferentially targets the region where the goal is located (Wilcoxon signed-rank test for Sector 1 vs. Sector 2, W = 32.5, p = 0.021, n = 9 sessions, one-sided). *p < 0.05. (D) Proportion of cells that maintained stable Distance-to-Goal tuning in both light and dark conditions (LD-Stable DTG: 43.5%).

Although RSC is closely associated with the visual system, lesions to RSC result in impaired navigation specifically under dark conditions[18, 49]. To investigate the role of RSC in navigation without visual information, we analyzed goal-directed navigation task performed in the dark (cue removed). Animals’ performance showed a slight decline (1.94±0.72 rewards/min) but remained adequate for navigation (Figure S8B). Despite the diminished visual information, we found that HD cells in RSC still stably maintained a preference for the direction of the original cue (Rayleigh p=0.01, n=71 cells; Figure 6A bottom), consistent with that under light condition.

Regarding distance perception, we found that 43.5% of the DTG neurons maintained a similar Distance-to-Goal tuning property across both light and dark conditions (Figures 6B and 6D). Furthermore, the population as a whole continued to encode Distance-to-Goal information, although with a decline compared to the light condition, and still exhibited a preference for the goal region. (Goal-Dark: Wilcoxon signed-rank test for Sector 1 vs. Sector 2, W=32.5, p=0.021, n=9 sessions, one-sided; Figures 6C, S8D and S8E), suggesting that, in the dark, some RSC neurons can retain their response patterns for goal-directed navigation that is typically seen under light conditions. These results are consistent with the possibility that RSC spatial representations are partly maintained by internal and self-motion-related signals when visual input is reduced.

## Discussion

In this study, we found that neurons in RSC encode the Euclidean distance between the animal and its goal during goal-directed navigation. Furthermore, we demonstrate the mixed-selective encoding pattern, as well as the population coding characteristics of distance-related signals within RSC. Subsequently, we compared the representation patterns of RSC neurons during goal-directed versus random foraging tasks, as well as under varying lighting conditions. These comparisons revealed that navigation selectively enhances allocentric head-direction encoding while leaving egocentric representations unchanged, and that these task-related spatial representations are maintained even when visual information is diminished.

However, the representation of distance does not exist only in RSC. In MEC, Butler et al. suggested that the firing patterns of neural populations, such as grid cells, are influenced by the goal and carry information about the distance between the animal and the goal[7]. In PPC, Alex et al. trained animals to pursue targets and found that PPC neurons represent moving targets based on the animal’s position, with the representation correlating to the target’s distance[8]. Additionally, Sarel et al. identified that in CA1 of bats, a subset of neurons encodes either the path distance or the Euclidean distance to the goal area[6]. Similar research also reported that in rats navigating to the unmarked goals within virtual reality (VR) environment, a portion of CA1 neurons demonstrates tuning properties associated with path distance[50]. These studies suggest that neuronal populations across different brain regions are capable of encoding information related to goal distance. Behind the widespread presence of distance information in these brain regions, there may be a systematic computational network that enables animals to efficiently use this information to guide their navigation behavior.

Both theoretical and experimental research suggest that RSC may be involved in the transformation of information between allocentric and egocentric reference frames in the brain[23, 24, 37, 51–53]. Distance, as a shared parameter between the egocentric and allocentric coordinate systems, is specifically used by RSC to encode goal location in goal-directed task, which may reflect this involvement. Our study confirms that, during navigation task,

Distance-to-Goal is one of the most prominent representations in RSC. Moreover, the population of cells involved in encoding this information has been found to contain distinct subpopulations encoding allocentric and egocentric information. We hypothesize that RSC may form a representation of Distance-to-Goal by integrating top-down distance information from the medial temporal lobe and bottom-up egocentric goal information from PPC. Future experiments utilizing multi-area simultaneous recordings could provide deeper insights into the dynamic processes and underlying differences in how distance information is transmitted and encoded across these pathways.

The updated separability–smoothness analysis refines the interpretation of mixed selectivity in RSC. The same population can encode multiple task variables while organizing those variables with different geometric emphases. In particular, Distance-to-Goal showed relatively high local smoothness, whereas egocentric boundary bearing showed stronger macro-scale separability. This pattern is consistent with RSC embedding goal-distance information in a locally coherent population state while maintaining a globally discriminable egocentric boundary reference. Such an organization may help reconcile reports that RSC participates in both spatial reference-frame transformations and goal-directed navigation[10, 23, 45].

This interpretation should be treated as a population-level geometric description rather than a claim about a specific downstream readout or circuit mechanism. The separability– smoothness slope summarizes how each variable is arranged in the recorded embedding; it does not by itself establish discrete coding classes, optimal representational trade-offs, or causal use by downstream areas.

In a VR environment, RSC neurons have been found to encode visual landmarks, and task engagement enhance this representation[41]. Despite the differences between one-dimensional VR and freely moving environments, we observed similar phenomena in freely moving animals. In our goal-directed navigation task, the preferred tuning angle of HD cells are significantly concentrated in the direction of the landmark, with a stronger HD signal representation compared to the random foraging task. Notably, we did not find much influence from the goal on the preferred tuning angles of HD cells, suggesting that the observed phenomenon is due to the animals’ navigational demands enhancing landmark-based spatial representation, rather than being related to the specific location of the goal. It is important to note that, since our visual landmark is a cue located on the boundary of the maze, the HD tuning curves of cells specifically firing to the landmark can also pass the selection criteria. Therefore, in our analysis, we did not further distinguish landmark cells based on HD cell properties.

The brain’s spatial representation has been shown to maintain relative stability through path integration and intrinsic cues in the reduction of visual information, yet it still experiences gradual drift over time[54, 55]. In our goal-directed navigation task, RSC neurons are also influenced by lighting conditions. However, when visual information is reduced, the stability of their representation patterns is better maintained compared to the random foraging task: HD signals continue to be encoded with high intensity, remaining anchored to the original landmarks, while the representation of DTG is still present. Furthermore, the tuning patterns of some DTG cells remain consistent with those observed under normal lighting conditions. These findings suggest that RSC representations are not solely driven by immediate visual input and may incorporate memory-related or self-motion-related components.

In contrast to the selective enhancement of allocentric head-direction signals, egocentric representations in RSC were not strengthened during goal-directed navigation, and the decoding accuracy of egocentric boundary bearing was similar between goal-directed and random foraging tasks. To further examine whether egocentric coding contributes to goal-specific spatial representation, we identified the spatial location where each neuron’s egocentric tuning was strongest (MVL_max) and found that neurons with high MVL_max were distributed across the maze without concentration near the goal. Moreover, the overlap between MVL_max neurons and distance-encoding neurons was minimal, confirming that these represent largely distinct subpopulations. These results indicate that, in our task context, egocentric coding did not exhibit a preferential representation toward the goal.

Unlike the preference for egocentric coding of goal locations observed by Wang et al. in the lateral entorhinal cortex[9], we did not observe a comparable egocentric goal preference in RSC. Similarly, LaChance et al. found that neither postrhinal cortex nor medial entorhinal cortex exhibited goal-zone overrepresentation during hidden-goal navigation[56], with their referencepoint analysis showing no goal-centered clustering—closely paralleling our MVL_max findings. However, this does not imply that RSC egocentric signals are generally uninvolved in goal representation. Park et al. showed that egocentric vertex coding in granular RSC can be selectively enhanced at goal-located vertices[46], but their task employed polygonal environments with salient geometric anchors, whereas ours used a circular maze without geometric features. Additionally, our recordings were predominantly from dysgranular RSC, whereas theirs targeted granular RSC exclusively, and differences in subregion composition may also contribute to the divergent results. Future studies comparing egocentric coding across RSC subregions and task geometries will help clarify these distinctions.

We propose that RSC plays a crucial role in organizing goal-relevant spatial representations during navigation in two ways. First, during task engagement, RSC encodes distance-to-goal information through a functionally coherent subpopulation that is selectively biased toward the goal, while simultaneously enhancing allocentric head-direction signals anchored to environmental landmarks. Second, when visual information is reduced, RSC maintains these task-related representations, consistent with contributions from prior experience and selfmotion signals during goal-directed navigation. Beyond these task-dependent modulations, we found that co-encoded behavioral variables in RSC occupy distinct positions in separability– smoothness space, suggesting that RSC organizes navigation-relevant information not only by content but also by measurable population geometry. Overall, these results suggest that RSC encodes goal-relevant spatial information, adapts its representational strength according to task demands, and sustains these representations when external sensory cues are unavailable.

## Resource availability

## Acknowledgements

We thank Drs Jozsef Csicsvari and Chen Wang for critically reading the manuscript. Our gratitude also goes to Yue Ke and Wei Wang for technical assistance. This work was supported by grants to H.X. from Brain Science and Brain-like Intelligence Technology - National Science and Technology Major Project (2022ZD0205000), the National Natural Science Foundation of China (32171038 and W2433058), and Guangdong Basic and Applied Basic Research Foundation (2022A515011896 and 2024A1515011033)

## Author Contributions

H.X. conceived and supervised the project. Y.C. and L.T. set up the recording equipment. Y.C. performed the surgery. Y.C. and X.W. conducted the recording and data analysis. X.W. performed the population analysis. Y.C., X.W., and H.X. wrote the manuscript.

## Lead contact

Further information and requests for resources should be directed to and will be fulfilled by the lead contact, Haibing Xu.

## Materials availability

This study did not generate new unique reagents.

## Data and code availability

The electrophysiological, behavioral, and processed data reported in this paper are available from the lead contact upon reasonable request. Custom code used for analysis is available from the lead contact upon reasonable request.

## Materials and methods

### Experimental model and study participant details

Five adult male Long-Evans rats (3–6 months old; 300–600 g) were used in this study. Prior to surgery, animals were group-housed under standard laboratory conditions (22 ± 1*^◦^*C, 50–60% humidity) with free access to food and water. Postoperatively, rats were individually housed in custom acrylic cages (51 *×* 37 *×* 32 cm^3^) under a 12-hour light/dark cycle.

During behavioral training and recordings, animals were maintained on a controlled diet to stabilize body weight at 85% of their baseline free-feeding weight, with water available at all times. All experimental protocols were approved by the Animal Care and Use Committee of Southern Medical University.

### Surgical procedures

All animals underwent surgical implantation of a custom 64-channel adjustable microdrive targeting the right hemisphere RSC. The microdrive comprised 16 tetrodes fabricated from 12-*µ*m tungsten wire, twisted and gold-plated to achieve tip impedance of approximately 300 kΩ. Surgeries were performed under isoflurane anesthesia (induction: 5%; maintenance: 1– 2%) with continuous physiological monitoring. Anesthesia depth was confirmed by the loss of toe-pinch reflex, after which animals were secured in a stereotaxic frame. The surgical site was prepared by shaving fur and disinfecting the scalp, followed by subcutaneous administration of saline (1 mL/h) to maintain hydration.

A midline scalp incision exposed the skull surface, followed by placement of two cerebellar grounding screws (for electrical reference) and 5–8 anchoring screws to secure the microdrive assembly. A craniotomy was drilled over the right hemisphere RSC using stereotaxic coordinates relative to bregma: −3.0 to −7.0 mm (anterior/posterior), 0–1.8 mm (medial/lateral).

The dura mater was carefully excised, and the microdrive was implanted under direct visual guidance. The cranial window was sealed with sterile petroleum jelly, and the assembly was affixed to the skull with dental cement. Postoperative care included analgesia (ibuprofen, 5 mg/kg) and antibiotic prophylaxis (enrofloxacin, 5 mg/kg), with ad libitum food access and individual housing. Animals recovered for at least seven days prior to experimental recordings.

### Histology

At the end of all experiments, the rats were anesthetized with isoflurane. An electrical lesion was then applied to each recording site using an Intan stimulator, with a 30 *µ*A current for 3 seconds, to facilitate the identification of the final recording sites. After 24 hours, the rats were perfused with physiological saline followed by 4% paraformaldehyde solution. Following perfusion, the brain tissue, along with the microdrive, was immersed in 4% paraformaldehyde for 24 hours to ensure thorough fixation.

Next, the electrodes were raised to their highest position and detached from the brain tissue to leave distinct electrode track marks in the tissue. After the electrodes were removed, the brain tissue was placed in a 30% sucrose solution for dehydration. Three days later, the brain tissue was sectioned into 40 *µ*m coronal slices and stained with Nissl stain. Finally, Brainrender was used to reconstruct the final recording sites.

### Behavioral task

Goal-directed navigation task: Briefly, the animals were trained in a cylindrical maze (diameter: 100 cm) with prominent visual cues on the wall (height: 50 cm). The animals were trained to locate an invisible circular goal zone (diameter: 20 cm) and remain there for at least 2 seconds, at which point an overhead food dispenser was activated to release a 45 mg pellet. Subsequently, the animals would leave the goal zone to search for and consume the reward. After that, the animals would continue navigating back to the goal zone to activate the next trial. The location of the goal zone remained consistent within each recording session, but varied across different days to prevent the animals from developing a long-term dependence on a specific location. Prior to surgery, the animals underwent training to ensure they could successfully perform the task post-implantation. Each recording session lasted either 20 or 30 min.

Random foraging task (RF): In the Random foraging task, the animals were placed in a different maze from the goal-directed navigation task (square: 100 cm *×* 100 cm *×* 40 cm or circular: diameter 100 cm, height 50 cm). Pellets were randomly scattered throughout the maze to encourage the animals to explore the entire environment. The recording duration lasted 20 min.

Dark recording: In the dark recording condition, all visible lights in the recording room were turned off, and indicator lights were covered with aluminum foil to ensure a completely dark environment. Infrared lights, placed above the maze, were used for illumination (invisible to the animals), to minimize visual information while still allowing the camera to record the animal’s behavior.

It should be noted that in some sessions, the RF task and goal-directed navigation task were recorded over two consecutive days without moving the tetrodes. Although we combined the recordings from both days for spike sorting, long-duration in vivo electrophysiological recordings make it challenging to reliably track individual neurons. Therefore, rather than examining the transition of single neurons between the two tasks, we focused on analyzing the proportion of specific cell types between tasks.

### Data acquisition equipment

Electrophysiological signals were recorded and digitized using an RHD2000 data acquisition system with a sampling frequency of 20 kHz. The microdrive implanted on each rat’s head was connected to the data acquisition system via an ultrathin SPI cable, and the signals were amplified using an RHD head stage (Intan Technologies). Behavioral data were recorded with a camera positioned above the maze at 50 Hz or 120 Hz (Basler) and synchronized with the electrophysiological signals via TTL. To facilitate subsequent analysis, the 50 Hz behavioral data were upsampled to 120 Hz.

### Behavioral variable extraction

For body part tracking, we used DeepLabCut. Specifically, we labeled 2,400 frames taken from five animals, of which 95% were used for training. We used a ResNet-50-based neural network with default parameters for 1,000,000 training iterations. We validated with one shuffle, and found the test error was 2.36 pixels and the training error was 2.25 pixels (image size was 1024 *×* 768 pixels). This network was then used to analyze videos from similar experimental settings.

Speed: We computed the animal’s speed using the central difference method. Specifically, for a given time point t, speed was calculated by subtracting the animal’s position at t - 10 frames from the position at t + 10 frames and dividing the difference by the corresponding time interval. The resulting speed was then smoothed using a Gaussian filter with a 0.4-second window. The smoothed speed was considered the animal’s instantaneous speed.

Head direction: The head direction was calculated as the rotation angle between the animal’s head vector and the maze’s x-axis, with an angle range of −180 to 180*^◦^*. A value of 0*^◦^* indicates that the animal’s head is aligned with the x-axis.

Distance-to-Goal: Distance-to-Goal is calculated as the Euclidean distance between the animal’s head position at each time point and the center of the goal zone. When the Distance-to-Goal is less than 10 cm, it indicates that the animal is within the goal zone.

Egocentric boundary bearing (Bearingboundary): The calculation of the bearing boundary is based on previous work[23]. Specifically, the Bearingboundary is calculated by first constructing a vector from the animal’s head to the nearest wall, and then computing the angular difference between this vector and the animal’s head direction vector. This angular difference is defined as the Bearingboundary. It is important to note that this angle has a fixed conversion relationship with the Bearingcenter, such that Bearingboundary = 180*^◦^* + Bearingcenter.

Unless otherwise specified, all data analyses were conducted using data collected when the animal’s speed was greater than 5 cm/s and the animal was not in a clear waiting phase.

### Spike sorting

Electrophysiological data were preprocessed using SpikeInterface, which included the removal of damaged channels and bandpass filtering (300–6000 Hz). Spike sorting was then performed using MountainSort4 via SpikeInterface, with a detection threshold set at 5. Further manual curation was carried out using Phy software. Well-identified cells were required to exhibit clear waveform asymmetry, with no significant noise during the refractory period, and a firing rate greater than 0.3 Hz.

### Rate maps and tuning curves

Spatial rate map: Two-dimensional histograms of the trajectory and corresponding spike positions were computed, with bin sizes set to 4 cm *×* 4 cm. A Gaussian filter (sigma = 1) was then applied to smooth both the trajectory and spike position histograms. The spatial rate map was obtained by dividing the two histograms. For presentation purposes, the spatial rate maps were rotated to ensure the goal position was always on the right side. Unless otherwise specified, all analyses were based on the unrotated maps.

MVL map: We first divided the entire maze, as well as the area outside the maze, into a grid composed of 6 cm *×* 6 cm bins. For both the RF and goal-directed navigation task, we applied this grid division to a 120 cm *×* 120 cm area, with the center of the maze located at the center of the grid. We systematically searched for the location of the MVLmax by calculating the MVL value of the corresponding egocentric bearing tuning curve when the reference point was positioned at the center of each bin. The resulting matrix of MVL values, representing the MVL for the tuning curve of each bin, was referred to as the MVL map.

Directional tuning curve: The full range of angles was divided into 30 bins (each representing 12*^◦^*). For each cell, the time spent in each bin and the corresponding spike count were calculated. The firing rate for each bin was obtained by dividing the spike count by the time spent in the bin. The resulting directional tuning curve was then smoothed using a Gaussian filter with sigma = 1.

Speed tuning curve: Similar to the direction tuning curve, the speed tuning curve was computed with a range from 5 cm/s to 55 cm/s, divided into 25 bins, each representing a 2 cm/s increment. The resulting speed tuning curve was then smoothed using a Gaussian filter with sigma = 1.

Distance-to-Goal tuning curve: Similar to the above, the Distance-to-Goal tuning curve was computed with a range from 10 cm (the center of the goal area, with a radius of 10 cm) to 60 cm, divided into 20 bins, each representing a 2.5 cm increment. It is important to note that distances greater than 60 cm were not included in the analysis due to the sparse nature of trajectories in this range, which could lead to the occurrence of outliers.

Egocentric boundary rate map (EBR): Following the method described in the previous study23, EBR were computed similarly to 2D spatial rate maps but referenced to the animal’s position. For each frame, distances to the maze boundaries were calculated in 1-cm bins, along with angular bins from −180*^◦^* to 180*^◦^* in 3*^◦^* intervals relative to the animal’s head direction. The angular bins were referenced such that 0*^◦^* was the front, 90*^◦^* left, 180*^◦^*/-180*^◦^* rear, and −90*^◦^* right. To avoid singular values, the animal’s maximum receptive radius was set to 40 cm, given the maze radius of 50 cm. Interpolation was performed on a higher-resolution grid, with angular and distance bins increased by factors of 6 and 5, respectively. Smoothed maps were obtained by convolving the raw data with a 2D Gaussian kernel (sigma = [120/7, 40/7]).

### Generalized linear model with forward feature selection

To systematically evaluate the encoding of multiple behavioral variables by individual RSC neurons, we employed a Poisson generalized linear model (GLM) with L1 regularization, implemented using the nemos package. For each neuron, spike counts were computed in non-overlapping time bins of 1/60 s. Five candidate behavioral variables were considered: head direction (H), speed (S), distance-to-goal (D), egocentric boundary bearing (E), and position (P, modeled jointly as x and y coordinates). Each variable was projected onto a set of basis functions prior to fitting: cyclic B-spline bases with 10 basis functions were used for the circular variables H and E, standard B-spline bases with 10 basis functions for S, 15 basis functions for D, and 8 basis functions per spatial dimension for P.

A forward feature selection procedure was used to determine the optimal combination of variables for each neuron. Starting from a null (intercept-only) model, candidate variables were iteratively added one at a time. At each step, the model incorporating each remaining candidate was evaluated via 10-fold cross-validation, and the information-theoretic loglikelihood gain (Δllh, in bits/spike) was calculated for every fold by comparing the candidate model to the null (uniform Poisson) model. A candidate variable was accepted only if it met two criteria: the mean Δllh gain across folds exceeded 0.01 bits/spike, and a one-sided Wilcoxon signed-rank test across folds yielded p < 0.05. The search terminated when no remaining variable satisfied both criteria.

### Measure used for cell type classification

For each recorded neuron, functional cell type was determined using a two-stage procedure. First, a classical tuning-based criterion was applied to identify candidate significant cells. Second, the results were intersected with the GLM forward search (see Generalized linear model with forward feature selection) to yield the final cell type assignments used in all analyses and figures.

Speed cell: The instantaneous firing rate for each cell was calculated with a 16.6 ms bin and a 250 ms smoothing window. The Pearson correlation coefficient between the instantaneous firing rate and the concurrent speed was computed as the cell’s speed score. A cell was considered a classical speed cell if its speed score exceeded the 99th percentile of the region-shuffle distribution. The final speed cell population was defined as the intersection of these classical speed cells with neurons whose GLM final model included the speed variable (S).

HD cell: For each cell, the mean vector length (MVL) of the head direction tuning curve was calculated as a measure of directional selectivity. A cell was considered a classical HD cell if its MVL exceeded the 99th percentile of the region-shuffle distribution. The final HD cell population was defined as the intersection of these classical HD cells with neurons whose GLM final model included the head direction variable (H).

DTG cell: For each cell, the Distance-to-Goal tuning curve was computed and compared with the distribution obtained from 1,000 circular shuffles. A DTG score was calculated as the maximum number of consecutive bins in the tuning curve that exceeded the shuffle-derived confidence interval. A cell was considered a classical DTG cell if its DTG score (max_point) was greater than or equal to 2.0 and its within-session tuning stability (Pearson correlation between the first and second halves of the session) was greater than or equal to 0.3. The final DTG cell population was defined as the intersection of these classical DTG cells with neurons whose GLM final model included the distance-to-goal variable (D).

EGO cell: The tuning curve for egocentric boundary bearing was computed and quantified using the MVL. A cell was considered a classical EGO cell if its MVL exceeded the 99th percentile of the region-shuffle distribution. The final EGO cell population was defined as the intersection of these classical EGO cells with neurons whose GLM final model included the egocentric variable (E).

Shuffling: For each cell, a circular shuffle of the entire spike train was performed by randomly shifting it within the range of [20 s, total length − 20 s], and this process was repeated 1,000 times to generate the shuffled dataset for that cell. The shuffled datasets from all cells recorded within the same brain region were combined to create the region-shuffle distribution, which served as the null distribution for the classical significance thresholds described above.

Note: The position variable (P) was included in the GLM to account for spatial modulation of firing rates but was not used to define an independent cell type; spatial firing properties such as spatial information and spatial stability were instead analyzed as continuous metrics to characterize DTG cells relative to other RSC neurons and simulated place cells (see Spatial rate map metrics).

### Spatial rate map metrics

The spatial information index was calculated following the function:

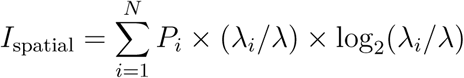

where *P_i_* is the occupancy probability of bin i in the firing rate map, *λ_i_* is the firing rate of bin i and *λ* the mean firing rate of the neuron.

To measure spatial coherence, we compute it using the unsmoothed firing rate map. For each bin in the firing rate map, we calculate the average firing rate of its neighboring bins within a 3×3 window, excluding NaNs. This produces a new map of neighbor rates. Spatial coherence is then calculated by determining the Pearson correlation between the original firing rate map and the neighbor rate map across the entire map.

### Tuning stability

For each cell, the stability of the rate map or tuning curve was defined as the Pearson correlation coefficient between the first and second halves of the recording session.

### Tuning curve fitting

For the Distance-to-Goal tuning curve of each cell, we used a linear model to fit the data in order to obtain the slope of the Distance-to-Goal tuning curve for each cell. The linear model is defined as:

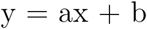

where a represents the slope, x is the corresponding Distance-to-Goal value, y is the firing rate at that point on the tuning curve, and b is the intercept of the fit.

### Neuron simulation

Egocentric Boundary cell: Using EBR, we quantify each real cell’s preferred angle and its dispersion (*σ*_angle), preferred distance and its dispersion (*σ*_distance), as well as the maximum and minimum firing rates. These parameters serve as inputs to the BoundaryVectorCells class in RatInABox, facilitating the development of a policy to simulate egocentric boundary cells.

Head Direction cell: From head direction tuning curves, we determine each real head direction cell’s preferred angle and angular spread (angular_spread_degree), alongside their maximum and minimum firing rates. These metrics are input into the HeadDirectionCells class provided by RatInABox to construct a policy for simulating head direction cells.

Speed cell: Utilizing the SpeedCell class in RatInABox, we derive the speed tuning range based on the mean speed and its standard deviation from actual behavioral data. The only required inputs are the maximum and minimum firing rates extracted from the real cell’s speed tuning curve.

Place cell: In the absence of place cell recordings within the target brain region, we generate 100 reference-free simulated place cells using the PlaceCells class in RatInABox. The place field radii are randomly assigned within 10 cm to 15 cm, and the maximum firing rate is uniformly set at 30 Hz.

Noise Addition and Cell Selection: For egocentric boundary, head direction, and speed cells, we introduce gradient noise by uniformly selecting five values within each cell’s firing rate range (defined by its maximum and minimum rates). We then compute the Pearson correlation between the tuning curves of the simulated and real cells at each noise level. The simulated cells exhibiting the highest correlation with their real counterparts are included in the simulated population. In contrast, place cells are generated by directly applying random noise at 0.1 Hz without subsequent cell selection.

Within RatInABox, agents are positioned in an environment matching the size of the experimental maze and execute behaviors derived from real animal trajectories. The simulation operates with a temporal resolution of 1/120 seconds, aligning with the experimental camera’s sampling rate. At each update step, the simulated firing rates are adjusted according to the policies of the various cell types. This iterative process ultimately produces a simulated cell population that closely mirrors the real population.

### Population dataset construction

For each session with N simultaneously recorded neurons, spike trains were divided into M time bins of 150 ms, followed by z-score normalization and smoothing with a Gaussian kernel (*σ* = 0.5), yielding an N *×* M matrix of population activity vectors. Simultaneously, the corresponding behavioral variables mentioned above were binned into 150 ms intervals, resulting in a one-dimensional matrix of length M. Both the population activity matrix and the label matrix served as inputs for subsequent supervised dimensionality reduction and decoding analyses.

### Dimensionality reduction

To perform dimensionality reduction and analyze the latent geometry of high-dimensional population activity, we employed CEBRA, a self-supervised representation-learning method for joint behavioral and neural data[57]. CEBRA generates consistent and high-performance latent spaces, enhancing the decoding accuracy of behavioral variables and ensuring robustness to domain shifts. The model was configured with the following hyperparameters: model_architecture=offset10-model, batch_size=64, learning_rate=0.01, temperature=1.0, output_dimension=3, max_iterations=5000, num_hidden_units=256, distance=cosine, conditional=delta_normal, verbose=True, and delta=0.3. All animal data were processed using these parameters for dimensionality reduction and latent space analysis.

### Macro-scale separability (structure index)

To quantify the global discriminability of behavioral-variable representations in the CEBRA-derived latent space, we computed the structure index (StrI), a kNN-based metric that measures how well data points belonging to different label bins are separated in embedding space[58, 59]. Continuous behavioral labels were first partitioned into B = 10 equally spaced bins based on the 5th–95th percentile range. For each pair of bins (a, b), the point clouds were pooled and k = 15 nearest neighbors were identified using Euclidean distance (accelerated via FAISS). The overlap between bins was defined as the fraction of each point’s k-nearest neighbors that belonged to the other bin, yielding a B *×* B asymmetric overlap matrix. From this matrix, the weighted out-degree of each bin was computed as the row-sum of overlaps, and the structure index was derived as

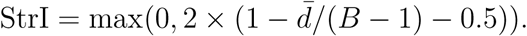

where *d̄* is the mean weighted out-degree across all bins. StrI ranges from 0 (complete overlap; random or uniform label distribution) to 1 (perfect separation; distinct, non-overlapping point clouds for each label bin). To assess statistical significance, a null distribution was generated by computing StrI on 100 label-shuffled permutations of the same embedding. StrI was computed independently for each of the five behavioral variables (Distance-to-Goal, Egocentric boundary bearing, Head direction, Position, Speed) on each cross-validation fold of the CEBRA embedding, and then averaged across folds.

### Micro-scale Smoothness (1*−* Normalized Local Neighborhood Consistency)

Local Neighborhood Consistency (LNC) was used to quantify local label inconsistency in the embedding. For each data point *x_i_* in the CEBRA latent embedding, the *k* = 30 nearest neighbors *N_k_*(*x_i_*) were identified using Euclidean distance, and the variance of the behavioral label within this neighborhood was computed:

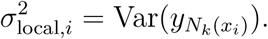

The raw LNC score was defined as the mean local variance across all points:

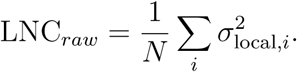

To enable comparison across variables with different label ranges, we normalized by the global label variance:

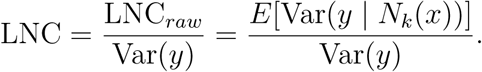

Lower normalized LNC indicates that nearby neural states carry more similar behavioral labels, whereas higher normalized LNC indicates weaker local label organization. For visualization and interpretation, we converted normalized LNC into Micro-scale Smoothness:

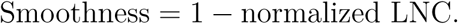

Thus, larger smoothness values indicate stronger local continuity of behavioral labels within the neural embedding. As with StrI, normalized LNC and Micro-scale Smoothness were computed for each behavioral variable on each cross-validation fold and averaged across folds.

### Representation Strategy quantification

To summarize the relative balance between macro-scale separability and micro-scale smoothness, each behavioral variable was represented in separability–smoothness space. For each variable, we estimated the principal trajectory slope across sessions in this two-dimensional space, where positive slopes indicate increasing smoothness with separability. The dashed line at *m* = 1 was used as a visual unity reference. For the stability analysis in Figure 4F, GoalLight and GoalDark sessions were pooled and bootstrap resampling (n = 10,000 iterations) was used to estimate the mean slope and 95% confidence interval for each variable. In each bootstrap iteration, data points were resampled with replacement, the principal trajectory was re-estimated, and the slope was recomputed. Pairwise slope differences were evaluated using permutation tests based on an ANCOVA model of smoothness as a function of StrI, variable identity, and their interaction. The observed reduction in residual sum of squares attributable to the interaction term was compared against a null distribution generated by permuting group labels, yielding a two-sided p-value.

### Explainable Boosting Machine (EBM) model configuration and training

We utilized the EBM from the interpret.glassbox package (https://github.com/interpretml/interpret) to decode neural activity and assess the relationship between neuronal population dynamics and various behavioral variables. The model was configured with key hyperparameters: learning_rate=0.1, max_bins=128, interaction_bins=32, n_estimators=300, and min_samples_leaf=10. This setup was carefully selected to balance interpretability and model complexity. Decoding was performed for each behavioral variable individually, and model performance was rigorously evaluated using ten-fold cross-validation. This framework enabled a detailed examination of how distinct neural firing patterns contribute to specific behavioral outcomes, ensuring robust and reliable insights into the neural-behavioral relationship.

### Feature selection based on Importance Scores

To identify neurons that most significantly contribute to predicting behavioral variables, we performed feature selection using importance scores from the EBM. In EBM, the importance score reflects how much each feature (in this case, neuronal activity) influences the model’s predictions. We conducted a ten-fold cross-validation, dividing the dataset into ten temporal subsets. For each fold, we computed the importance scores for all neurons and selected the top 50% of neurons with the highest scores. Neurons that were selected in at least eight out of ten folds were considered important, ensuring that the final selection was robust and reliable.

### Partial Dependence Plot (PDP) generation and Interpretation

To examine the relationship between neuronal firing rates and predicted behavioral responses, PDPs were generated for both important and non-important neurons. For each important neuron, firing rates were varied within the 0–90th percentile range, discretized into 20 equally spaced values. At each value, the neuron’s firing rate was set to the current value while keeping other features constant, and the EBM predicted the corresponding behavioral response. The PDP for each neuron was obtained by averaging these predictions, followed by smoothing with a Gaussian filter (sigma = 1) for clarity.

Quantitative metrics were derived from the PDPs, including the range of predicted responses, which indicates the span of behavioral outcomes; and the linear slope, which reflects the direction and strength of the relationship between firing rate and behavior.

In the PDP plots, important neurons were visualized with color-coding based on the sign of their linear slope green for positive slopes and blue for negative slopes with opacity proportional to the absolute value of the slopes to highlight neurons with stronger effects. Non-important neurons were represented as shaded grey areas, providing an overview of general response patterns without specific directional trends.

### Effect of cell number on decoding performance

To assess the impact of neural population size on decoding, we performed decoding on subgroups of 5, 10, 15, and 20 neurons (increasing by 5 neurons at each step). For each population size, we randomly sampled 100 times and calculated the mean decoding performance along with the 95% confidence intervals. The resulting plot illustrates how decoding performance varies as the neural population size increases.

### Attention map

The maze area of the animal’s activity was divided into 177 square grid units, each measuring 6.6 cm *×* 6.6 cm, with the center of each unit serving as a reference point. The realtime distance from the animal to each reference point was computed during the behavioral experiment. Using the recorded neural population activity, we decoded the distances to these reference points, obtaining decoding performance expressed as R-squared values for all grid units. This reflects the distribution of neural population encoding strength across different areas of the maze. To examine whether the animal’s distance encoding exhibits spatial preferences, we normalized the attention map using min-max scaling. The maze was then split into two halves along a line perpendicular to the line connecting the goal and the maze center, passing through the center. The halves were designated as sector 1 (containing the goal) and sector 2. The average decoding performance for each half (using the normalized R-squared values) was then calculated, and statistical differences were assessed across sessions using the Wilcoxon signed-rank test.

### Quantification and statistical analysis

Statistical analyses were performed using custom Python scripts with the scipy.stats library. All statistical tests were non-parametric, and data were not subjected to normality tests. For all box plots, the central mark indicates the median, and the bottom and top edges of the box represent the 25th and 75th percentiles, respectively. The whiskers extend to the most extreme data points within 1.5 times the interquartile range from the bottom or top of the box, and any data points beyond this range are plotted individually using a diamond symbol. Statistical details are provided in the main text or in figure legends.

## Supplementary Materials

**Figure S1.**
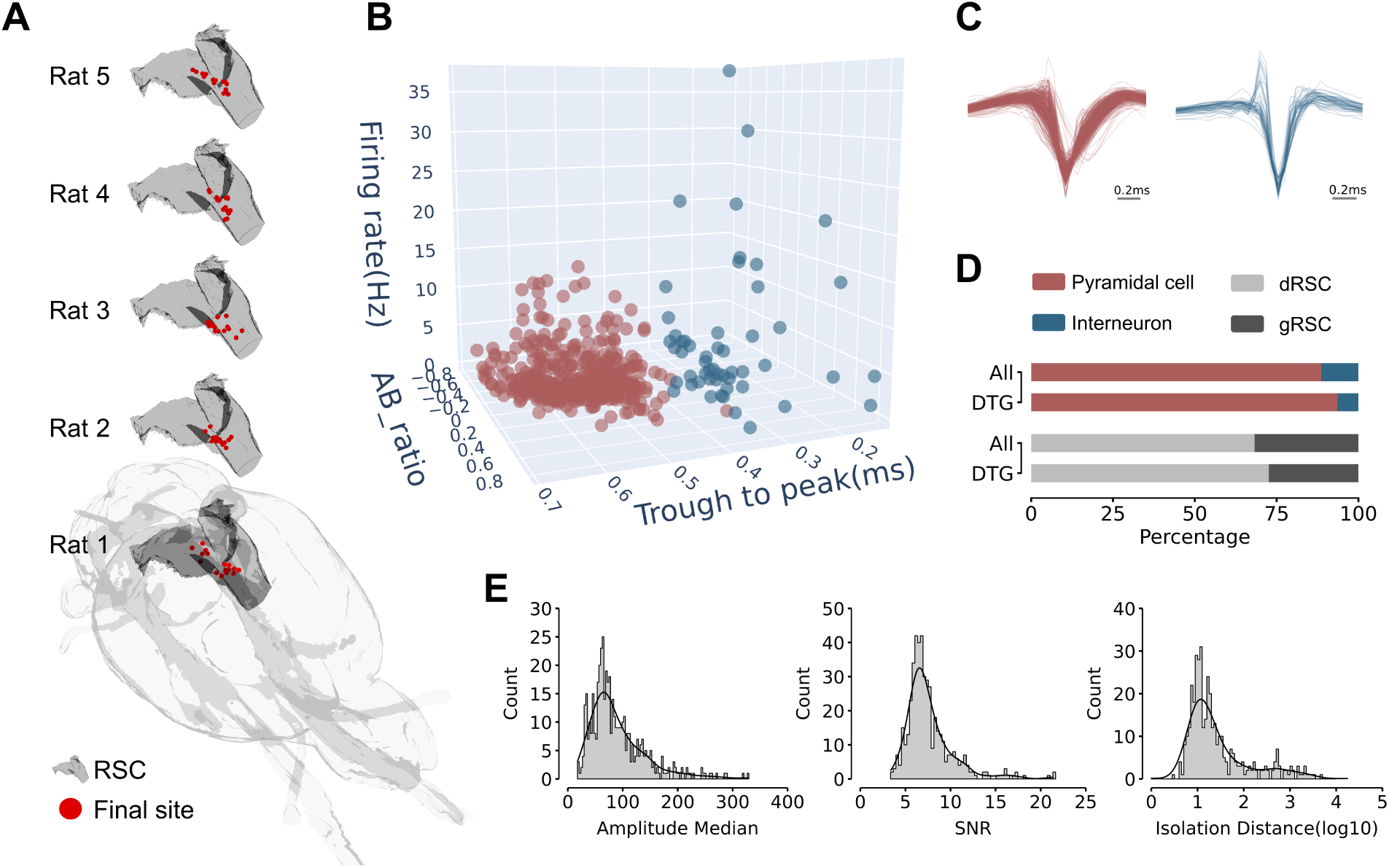
Classification of neurons and histological details, related to Figure 1. (A) Histological reconstruction of recording sites. For each animal, the final recording locations of all tetrodes (red dots) were reconstructed in RSC (dark gray). All recording sites were confirmed to be within RSC. (B) Neurons in RSC were classified into putative pyramidal neurons (trough-to-peak *≥* 0.4 ms, red dots) and putative interneurons (trough-to-peak *<* 0.4 ms, blue dots) based on trough-to-peak duration, firing rate, and waveform asymmetry (AB ratio), displayed in a three-dimensional feature space. (C) Representative waveforms of all putative pyramidal neurons (red) and putative interneurons (blue). (D) Proportion of DTG neurons based on neuronal type and recording location. Among DTG neurons, pyramidal neurons (DTG-Pyr: 93.5%, 58/62, red) and interneurons (DTG-Int: 6.5%, 4/62, blue) exhibited distributions consistent with those observed for all recorded cells (ALL-Pyr: 88.6%, 410/463; ALL-Int: 11.4%, 53/463). The distribution of DTG neurons across the dRSC (light gray) and the gRSC (dark gray) was as follows: DTG-dRSC: 72.6% (45/62) and DTG-gRSC: 27.4% (17/62), which was comparable to the distribution of all recorded cells (ALL-dRSC: 68.3%, 316/463; ALL-gRSC: 31.7%, 147/463). (E) Quality control metrics for spike sorting. From left to right, histograms illustrate the median amplitude, signal-to-noise ratio (SNR), and isolation distance of all neurons recorded during the goal-directed navigation task.

**Figure S2.**
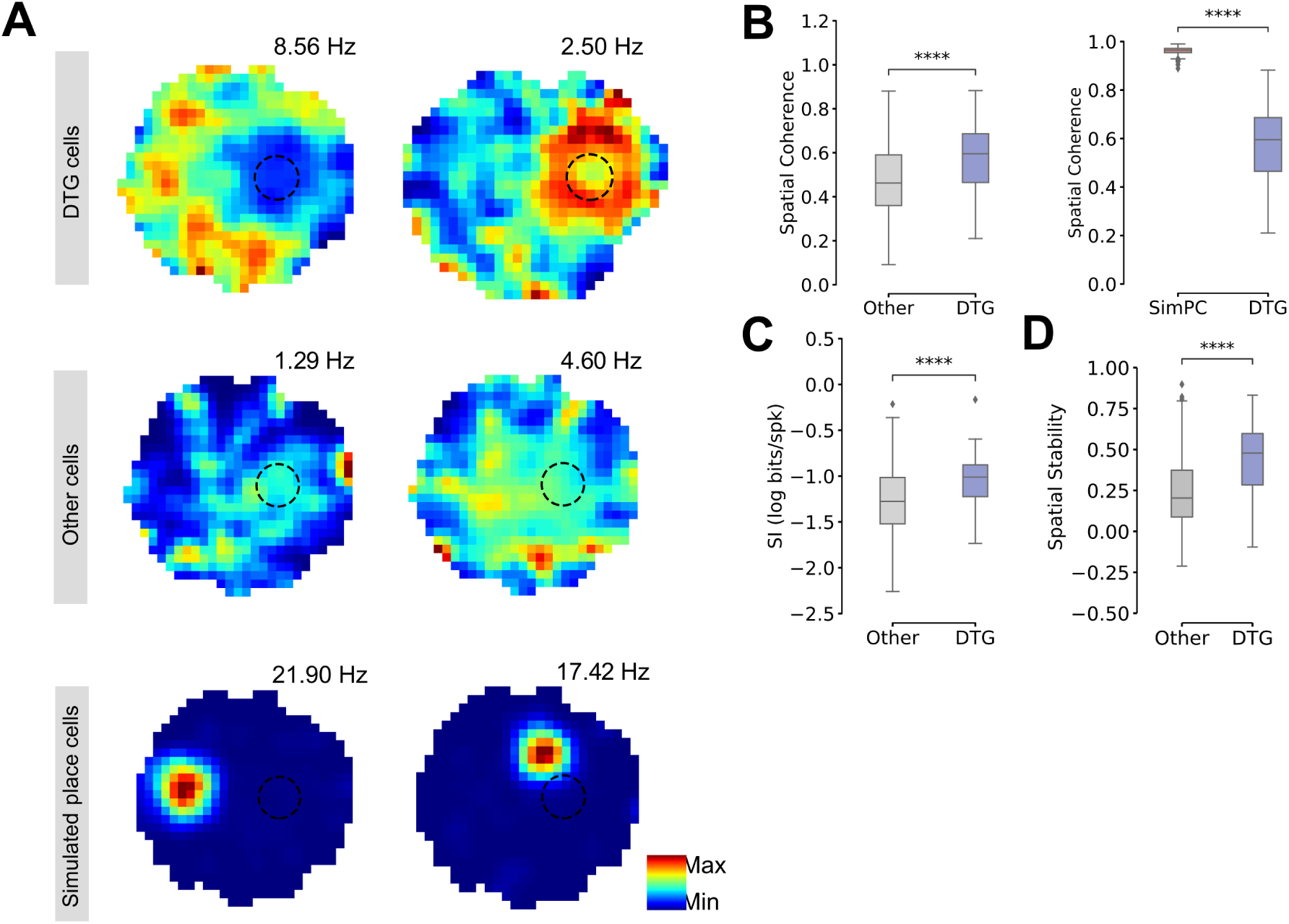
Comparison of DTG cells tuning, related to Figure 1. (A) Example firing rate maps for DTG cells, other RSC cells, and simulated place cells. (B) Comparison of spatial coherence. Left: Comparison between other RSC cells and DTG cells; Right: Comparison between simulated place cells and DTG cells. DTG cells exhibit higher spatial coherence than other RSC cells but lower coherence than simulated place cells. ****p < 0.0001, Wilcoxon Rank-Sum Test. (C) Comparison of spatial information between other RSC cells and DTG cells. DTG cells exhibit significantly higher spatial information than other RSC cells. ****p < 0.0001, Wilcoxon Rank-Sum Test. (D) Comparison of spatial stability between other RSC cells and DTG cells. DTG cells show greater spatial rate map stability than other RSC cells. ****p < 0.0001, Wilcoxon Rank-Sum Test.

**Figure S3.**
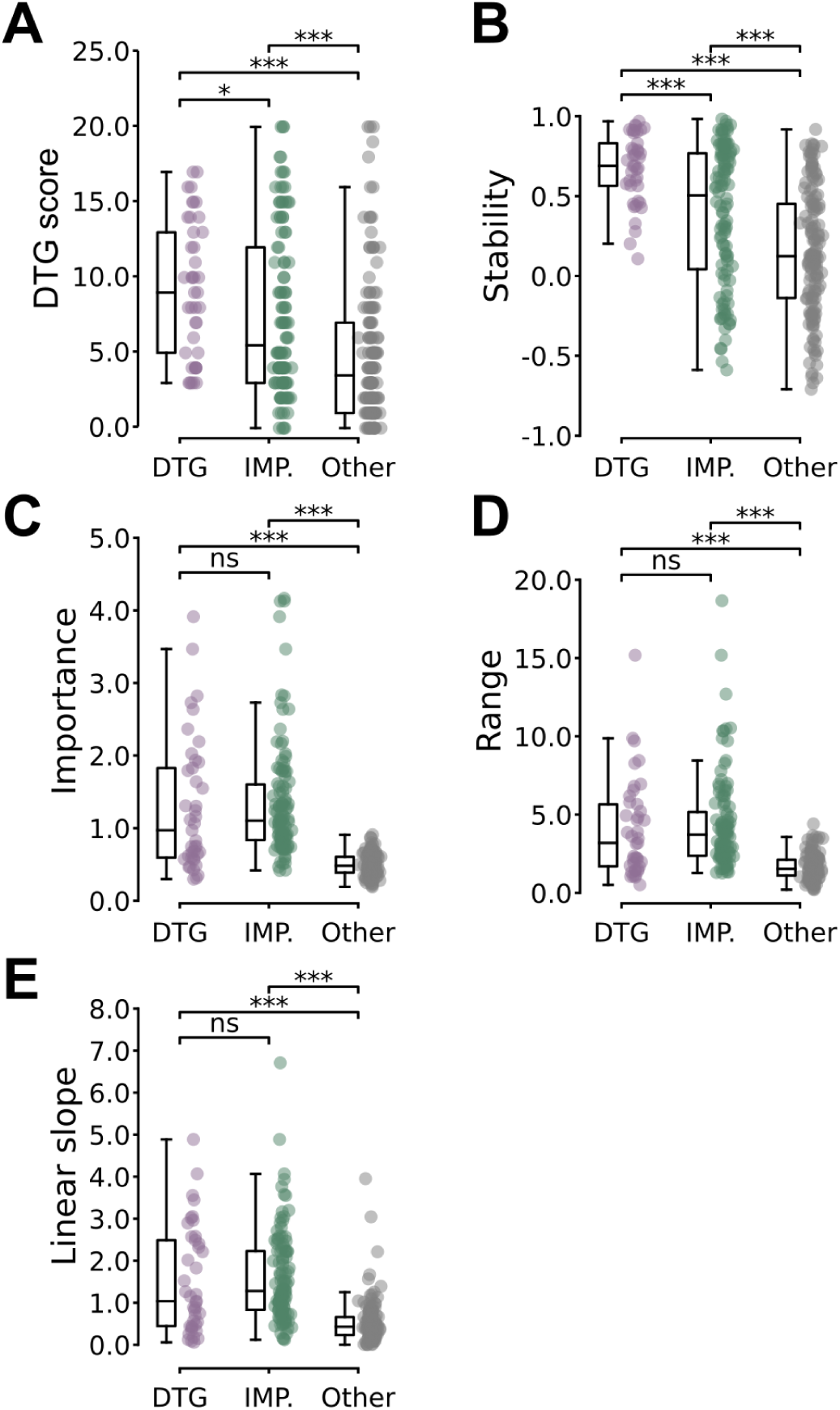
Comparison of Distance-to-Goal tuning properties across DTG, IMP, and Other cells, related to Figure 2. (A) DTG score – This metric quantifies the degree of Distance-to-Goal tuning in each neuron.Both DTG and IMP cells show significantly higher DTG scores compared to Other RSC cells, with DTG cells exhibiting the strongest tuning. ***p < 0.001, Mann-Whitney U test. (B) Stability – This measure reflects the stability of the tuning curve over time, with higher values indicating more consistent tuning. The statistical comparison reveals that DTG and IMP cells have significantly higher stability than Other RSC cells, indicating that DTG and IMP cells maintain a more consistent response profile in relation to the task. However, the stability of IMP cells is lower than that of DTG cells. ***p < 0.001, Mann-Whitney U test. (C) Importance – This measure quantifies the relative importance of neurons in predicting behavioral variables. The statistical analysis demonstrates that DTG and IMP cells exhibit significantly higher importance than Other RSC cells, indicating their stronger contribution to the prediction model. ***p < 0.001, ns > 0.05, Mann-Whitney U test. (D) Range – This parameter reflects the span of neural responses. The statistical comparison shows that DTG and IMP cells have a significantly broader range of responses compared to Other RSC cells, suggesting more dynamic and informative neural activity. ***p < 0.001, ns > 0.05, Mann-Whitney U test. (E) Linear slope – The linear slope indicates the directionality of the neural response. The statistical analysis reveals that DTG and IMP cells show significantly steeper slopes than Other RSC cells, highlighting their stronger directional effects in the model. ***p < 0.001, ns > 0.05, Mann-Whitney U test.

**Figure S4.**
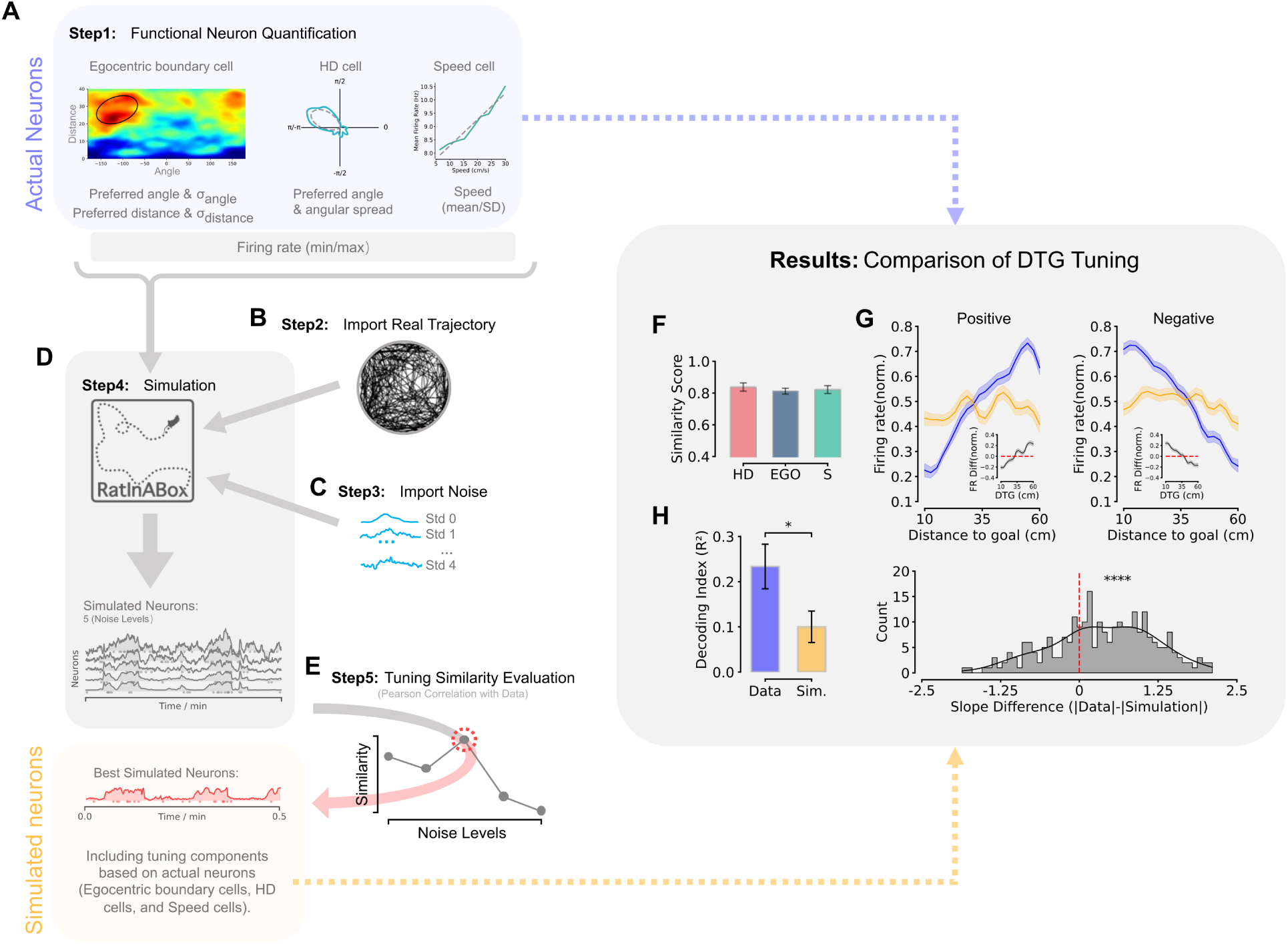
Workflow for simulating functional neurons and comparison of DTG tuning, related to Figure 3. (A) Step 1: Functional Neuron Quantification. For egocentric boundary cells, the preferred angle, preferred distance, angular spread (*σ_angle_*), and distance dispersion (*σ_distance_*) are quantified from the boundary rate maps. For head direction cells, the preferred angle and angular spread are determined from head direction tuning curves. The tuning characteristics of speed cells are determined based on the mean speed and standard deviation from the animal’s real trajectory. These parameters are used as inputs to the corresponding cell policies (BoundaryVectorCells, HeadDirectionCells, and SpeedCell) in RatInABox. (B) Step 2: Import Real Trajectory. Real animal trajectory data is imported to align the simulation with actual behaviour, providing the basis for agent positioning and movement in the virtual environment. (C) Step 3: Import Noise. Noise levels (Std 0, Std 1, and Std 4) are added to the firing rates of the simulated neurons, with five noise values selected within the range of each real cell’s firing rates to simulate different noise conditions. (D) Step 4: Simulation. RatInABox is used to simulate the activity of neurons in a virtual environment corresponding to the experimental maze. The simulation generates firing rate patterns for each cell type (egocentric boundary bearing, head direction, and speed cells) based on the defined policies and noise levels. (E) Step 5: Tuning Similarity Evaluation. The Pearson correlation between the tuning curves of simulated and real cells is calculated at each noise level. The best-simulated neurons are identified based on their highest correlation with real neurons, ensuring that the simulated population closely mirrors the real population for further analysis. (F) Comparison of tuning similarity between simulated and actual cells for three functional cell types (HD, EGO, Speed). Error bars represent mean *±* 95% CI of the data. (G) Comparison of Distance-to-Goal tuning between functional and simulated neuronal populations. Top left: Normalized firing rates showing a positive correlation with distance in functional cells (blue) and their corresponding simulated cells (orange). Inset: Difference in Distance-to-Goal tuning (functional minus simulated). Top right: Similar to the left, but depicting a negative correlation. Bottom: Distribution of slope differences (|Data*|−|*Simulation|) between functional and simulated cells. The positive shift indicates that functional cells exhibit significantly steeper Distance-to-Goal tuning slopes than their simulated counterparts. ****p < 0.0001. (H) The functional cell population significantly outperforms the corresponding simulated cells in Distance-to-Goal decoding. Error bars represent mean *±* SEM of the data. *p < 0.05.

**Figure S5.**
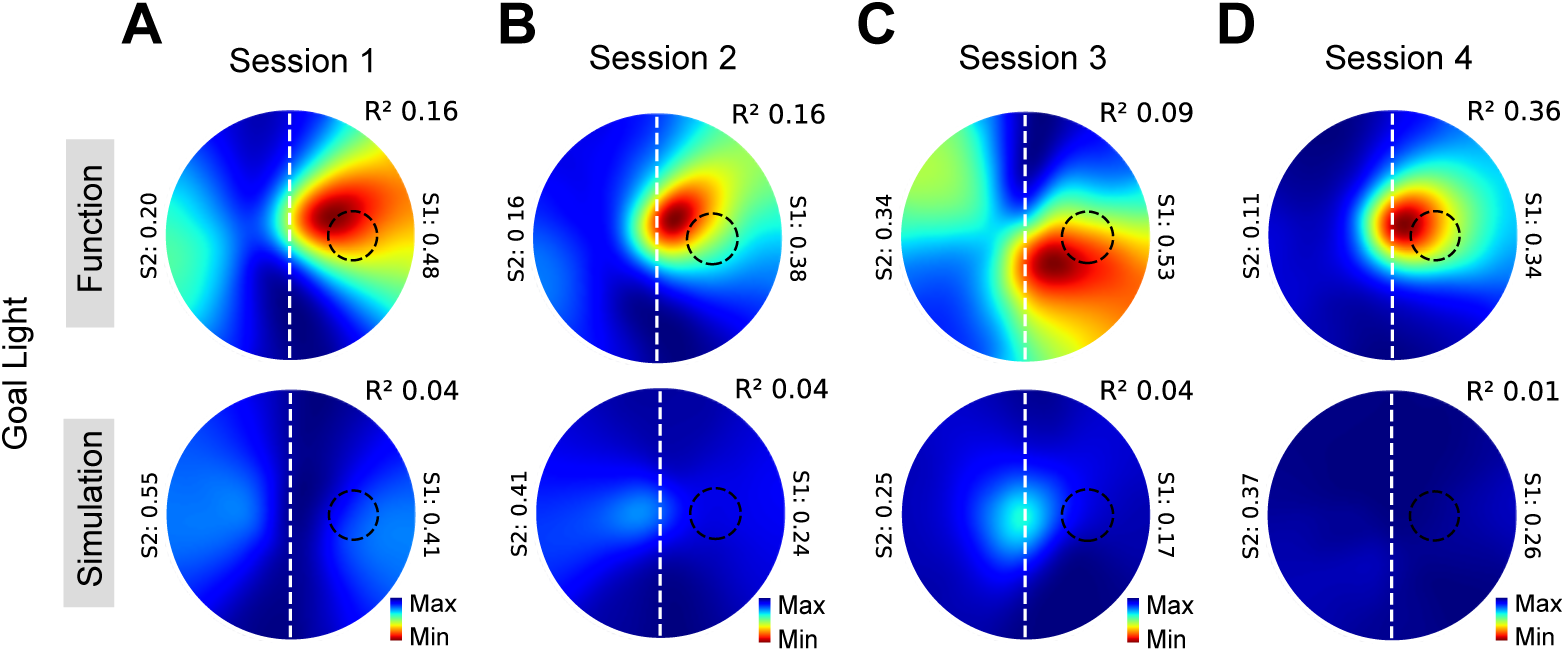
Attention maps for functional and simulated cells across sessions, related to Figure 3. (A–D) Attention maps showing the spatial preference of functional and simulated cells for four example sessions. Only the functional cells exhibit a goal-preference distance representation, while both populations share the same influence from goal-directed non-uniform trajectories, ruling out behavioral biases.

**Figure S6.**
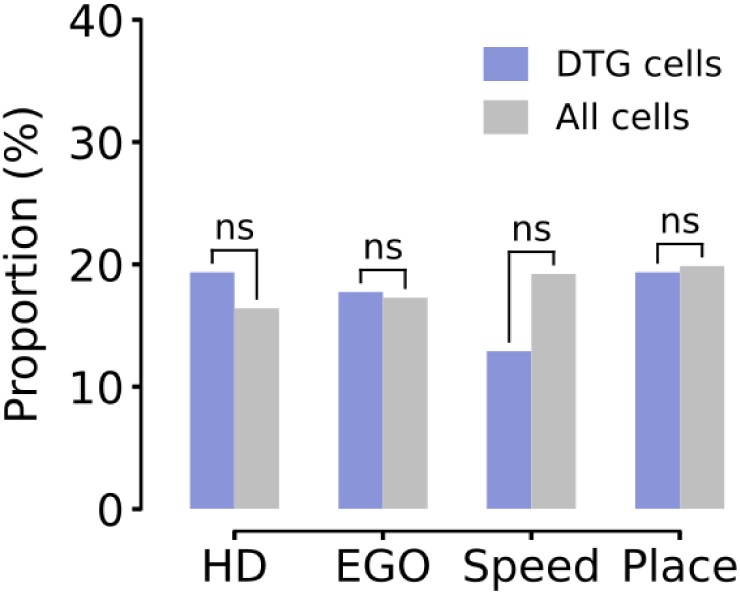
Functional cell type proportions within the DTG population, related to Figure 4. Comparison of the proportions of four functional cell types HD, EGO, Speed, and Place between DTG cells (blue) and the overall recorded population (gray). None of the four cell types was significantly enriched in the DTG population (HD: DTG = 19.4%, All = 16.4%, p = 0.63; EGO: DTG = 17.7%, All = 17.3%, p = 1.0; Speed: DTG = 12.9%, All = 19.2%, p = 0.24; Place: DTG = 19.4%, All = 19.9%, p = 1.0; Chi-squared test), indicating that Distance-to-Goal encoding is broadly distributed across functional cell types in RSC rather than being preferentially coupled with any single type of spatial or motor representation. ns > 0.05.

**Figure S7.**
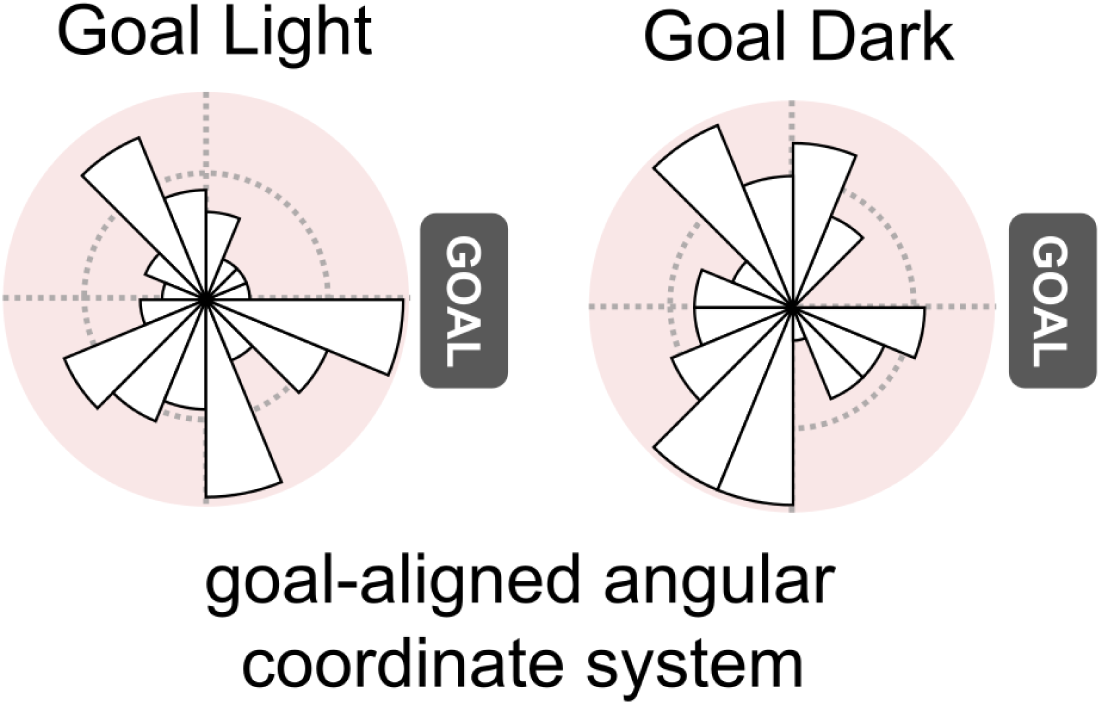
Further analysis of HD Cells, related to Figure 6. In the Goal-aligned angular coordinate system, we did not observe any asymmetric distribution of preferred angles for HD cells (Goal Light: p = 0.24; Goal Dark: p = 0.88; Rayleigh test).

**Figure S8.**
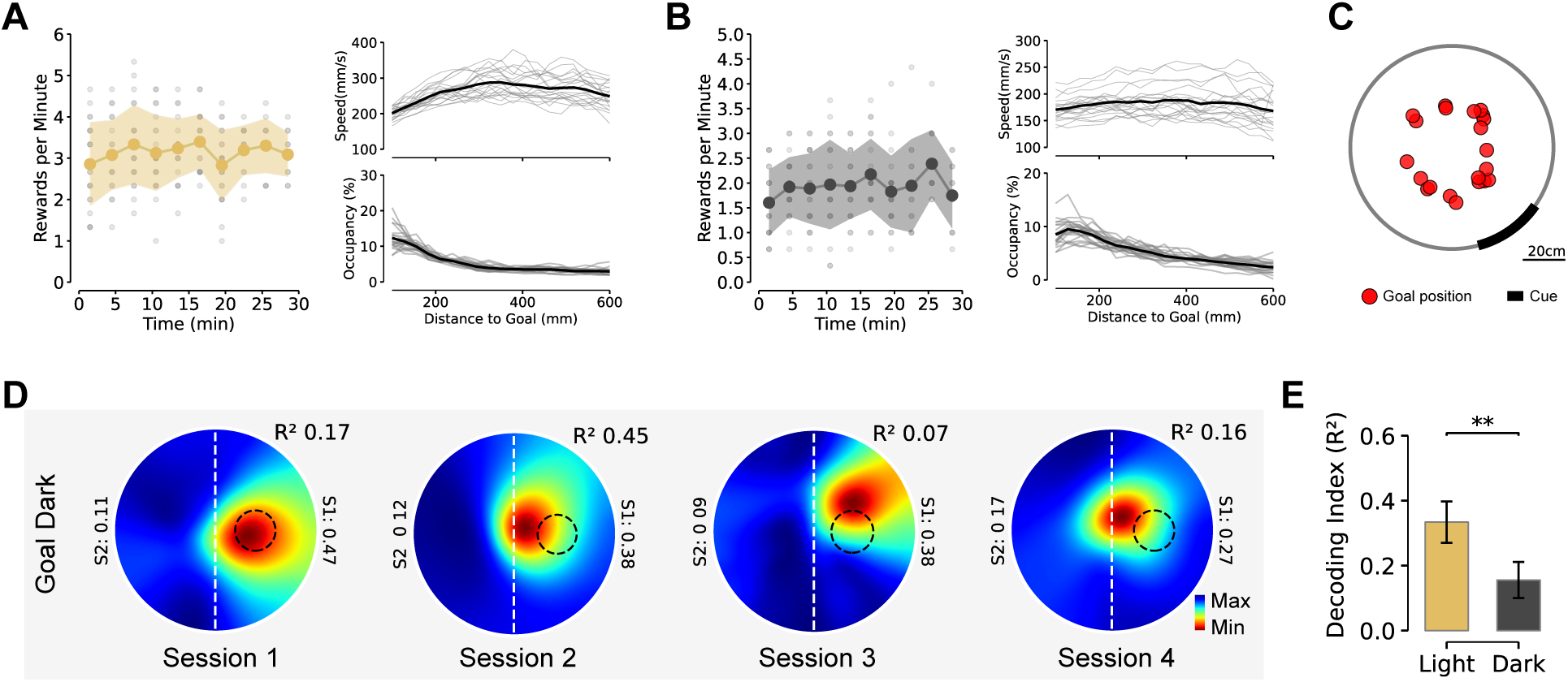
Behavioral quantification and decoding of goal navigation task under light and dark conditions, related to Figures 1 **and 6**. (A) Behavioral quantification of the goal-directed navigation task under light conditions. Left: Reward rate per minute (mean *±* SD, yellow) during the light condition, with gray representing data from individual sessions, as also shown in Figure 1C. Right: Relationship between animal speed, trajectory occupancy, and Distance-to-Goal, with gray lines representing data from each session and the black line indicating the average. (B) Similar to panel A, behavioral quantification of the goal-directed navigation task in the dark condition. The reward rate and trajectory occupancy indicate that animals are still able to perform the goal-directed navigation task with reduced visual information, although the reward rate per minute (mean *±* SD, 1.94 *±* 0.73) was lower than in the light condition. (C) Cue and goal zone center positions within the maze for each recording session. As the goal position varied across sessions, we could assess the relative influence of the goal and cue on animal behavior. (D) Attention maps from four example sessions of the goal-directed navigation task performed in the dark. (E) Comparison of Distance-to-Goal decoding performance under light versus dark conditions. While decoding accuracy for Distance-to-Goal was significantly lower in the dark condition, RSC population still encoded Distance-to-Goal information. **p < 0.01, Wilcoxon signed-rank test.

